# Loss of MC1R signaling implicates TBX3 in pheomelanogenesis and melanoma predisposition

**DOI:** 10.1101/2023.03.10.532018

**Authors:** H. Matthew Berns, Dawn E. Watkins-Chow, Sizhu Lu, Pakavarin Louphrasitthiphol, Tongwu Zhang, Kevin M. Brown, Pedro Moura-Alves, Colin R. Goding, William J. Pavan

## Abstract

The human Red Hair Color (RHC) trait is caused by increased pheomelanin (red-yellow) and reduced eumelanin (black-brown) pigment in skin and hair due to diminished melanocortin 1 receptor (MC1R) function. In addition, individuals harboring the RHC trait are predisposed to melanoma development. While *MC1R* variants have been established as causative of RHC and are a well-defined risk factor for melanoma, it remains unclear mechanistically why decreased MC1R signaling alters pigmentation and increases melanoma susceptibility. Here, we use single-cell RNA-sequencing (scRNA-seq) of melanocytes isolated from RHC mouse models to reveal a Pheomelanin Gene Signature (PGS) comprising genes implicated in melanogenesis and oncogenic transformation. We show that TBX3, a well-known anti-senescence transcription factor implicated in melanoma progression, is part of the PGS and binds both E-box and T-box elements to regulate genes associated with melanogenesis and senescence bypass. Our results provide key insights into mechanisms by which MC1R signaling regulates pigmentation and how individuals with the RHC phenotype are predisposed to melanoma.

## Introduction

Cancer predisposition is usually defined by inherited genetic mutations which increase an individual’s risk of developing the disease. Identification of the genes and molecular pathways underlying predisposition is essential for both cancer treatment and prevention. Melanoma, the most lethal form of cutaneous cancer, arises from melanin-producing melanocytes and represents an excellent model to study cancer risk factors. Melanoma is an increasingly common skin cancer caused by high levels of sun exposure and subsequent damage by ultraviolet (UV) light. All stages of the disease are amenable to study and the genetic basis of disease initiation is well-defined. Specifically, cutaneous melanoma usually arises from acquisition of activated pro-oncogenes (e.g., BRAF, NRAS) combined with senescence bypass through inactivation of cell cycle regulators (e.g., CDKN2A/INK4A)^1–4^. However, in addition to genes falling into these two categories, additional genetic loci have been implicated in melanoma predisposition. Understanding the molecular basis for melanoma predisposition is a key issue which has the potential to inform melanoma prevention strategies.

Red Hair Color (RHC) is a well-described melanoma predisposition phenotype. Individuals with the RHC phenotype, possessing red hair, pale skin, freckles, and an inability to tan, show an increased risk of developing melanoma compared to those with a non-RHC disposition^5–7^. The RHC trait is mostly explained by coding variants of reduced function in the *melanocortin 1 receptor (MC1R)* gene which encodes a G-protein-coupled receptor (GPCR) located on the cell surface of melanocytes. MC1R signaling regulates pigment/melanin production^7–9^; therefore, diminished MC1R function alters the ratio of cellular eumelanin and pheomelanin that largely determines skin, hair, and eye color. Decreased MC1R signaling results in the loss of photoprotective eumelanin synthesis and in turn allows for almost exclusive production of pheomelanin. High levels of pheomelanin in RHC individuals exacerbate DNA damage through the amplification of free radicals which is worsened when cells are exposed to UV^10,11^, and pheomelanin itself can contribute to melanomagenesis in the absence of UV radiation^11^. These observations may partly explain why individuals with genetic RHC variants in *MC1R* exhibit an increased somatic mutational burden in melanocytes^12^.

Although the influence of *MC1R* RHC variants on melanin composition may play a role in melanoma predisposition, *MC1R* mutation status may also have an impact on melanoma initiation unrelated to its role in pigmentation. For example, individuals with *MC1R* variants fail to activate PTEN upon UV exposure, resulting in increased PI3K/AKT signaling and oncogenic transformation^13^. However, whether MC1R signaling plays additional roles in the prevention of melanoma initiation remains unknown. In part, this may be because, by definition, a molecular mechanism underlying melanoma predisposition must already be operating prior to the acquisition of the ultimate trigger for melanoma initiation, such as acquiring activating BRAF or NRAS mutations.

MC1R signaling shows high evolutionary conservation among mammals, thus murine species provide an excellent model to explore this pathway. This includes mice with active MC1R signaling (nonagouti; *a/a*), and suppressed MC1R signaling (recessive yellow, *Mc1r^e/e^*, and lethal yellow, *A^y^/a*). Nonagouti mice have a dark coat color that results from a loss-of-function (LOF) allele of the nonagouti gene (*a*), which encodes the mouse homolog of agouti signaling protein (ASIP). In the absence of this important MC1R antagonist, nonagouti melanocytes predominantly synthesize eumelanin. In contrast, recessive yellow (*Mc1r^e/e^*) and lethal yellow (*A^y^/a*) mice provide two models for RHC, each with melanocytes that synthesize primarily pheomelanin due to distinct molecular mechanisms which cause reduced MC1R signaling. Recessive yellow mice carry a LOF allele in the MC1R receptor while the similar yellow coat color of lethal yellow mice is caused by constitutive overexpression of the MC1R antagonist, agouti signaling protein^14–16^.

Given the role of the RHC trait in melanoma risk, a better understanding of the genetic regulatory network underlying aberrant MC1R signaling is needed. Previously, April et al^17^ identified several novel genes linked to pheomelanogenesis in mouse models using bulk RNA-sequencing. Since this study used whole skin layers, of which melanocytes comprise only a small fraction of the total cell population, gene expression from other cell types such as keratinocytes, basal cells, and dendritic cells potentially masked the detection of melanocyte-intrinsic changes. Haltaufderhyde and Oancea^18^ performed bulk RNA-sequencing on human epidermal melanocytes to assess global gene expression changes between light and dark colored cells. However, the genotypes were unknown, and melanocytes were not exclusively producing one form of melanin over the other. While these previous studies established an important foundation for our understanding of pheomelanogenesis, additional, high-resolution genomic studies are needed to fully evaluate the intricate molecular networks regulating eumelanogenesis, pheomelanogenesis, and melanoma development.

To better understand the gene regulatory network associated with the RHC trait, we used single-cell RNA-sequencing (scRNA-seq) to compare melanocytes with both active and suppressed MC1R signaling. We defined a set of gene candidates, the Pheomelanin Gene Signature (PGS), implicated in regulating pheomelanogenesis and melanomagenesis. We further explored the chromatin occupancy of TBX3, a PGS- and melanoma-associated T-box family member. Our results provide key insights into how RHC trait variants may increase melanoma risk via altered regulation of PGS candidate genes downstream of MC1R.

## Results

### Single-cell RNA-sequencing of primary melanocytes identifies a Pheomelanin Gene Signature (PGS)

To better understand global gene expression changes downstream of MC1R signaling, we used single-cell RNA-sequencing (scRNA-seq) to define the transcriptomic profiles of primary mouse melanocytes which predominantly synthesize either pheomelanin (lethal yellow, *A^y^*; or recessive yellow, *Mc1r^e^*) or eumelanin (nonagouti, *a*). The dorsal skin was removed from post-natal day 3 (P3) animals along anatomically defined margins consisting of four incisions (Fig 1A): two transverse cuts, with one between each axilla, and the other between the anterior pelvises, and bilateral incisions at the midline from the axilla to the anterior pelvis. This tissue was then enzymatically digested into a single-cell solution. Melanocytes comprise ∼5% of cells within the skin; therefore, after dissection and digestion of tissue, a fluorescent-activated cell sorting (FACS) approach was used to enrich the pool of melanocytes for sequencing^19^. Positively selected cells were defined as c-KIT^+^ and CD45^-^ (Fig 1A,B). To eliminate selection bias among potential subpopulations of melanocytes that may vary in c-KIT levels, a lower intensity threshold for c-KIT positivity was used, yielding a population enriched for melanocytes but expected to also contain other cell types. Isolated cells were processed for library construction and RNA-sequencing via the 10x Genomics Chromium pipeline. After the implementation of quality control measures (see Methods), sequencing data was obtained from a total of 11,058 cells from all samples. The corresponding data underwent dimensional reduction via non-biased principal component (PCA) and graph-based clustering analyses for graphical display by Uniform Manifold Approximation and Projection (UMAP) (Fig 1C,D). We classified and labeled captured cell types by overlaying published murine skin cell gene signatures^20^ (Fig 1C,D; Table S1; Fig S1-S2). Cells classified as melanocytes included 4 of the 16 graph-based clusters (GBC) identified in cells isolated from a nonagouti and lethal yellow pair of samples, and 1 of 12 GBCs identified in cells isolated from a nonagouti and recessive yellow pair of samples. Subsequently, for each of the 2 pairwise experiments, all non-melanocyte cells were filtered out and only gene expression from the cells classified as melanocytes were used for further analysis.

**Figure 1.**
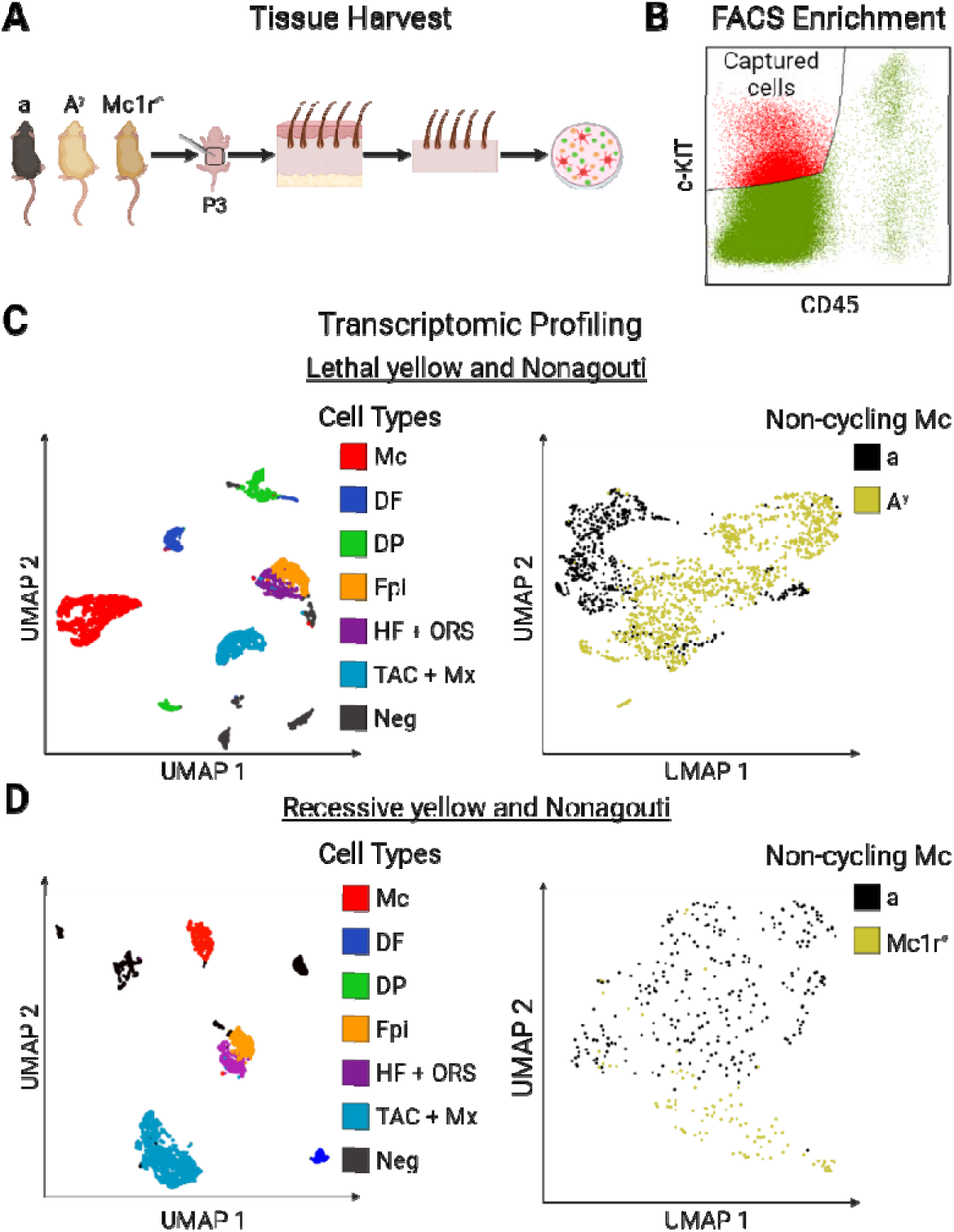
Overview of single cell sequencing of primary murine melanocytes. (A) Schematic of tissue harvest and digestion for melanocyte isolation from dorsal skin (a = nonagouti; A^y^ = lethal yellow, Mc1r^e^ = recessive yellow). (B) Representative FACS enrichment of c-KIT^+^/CD45^-^ cells isolated from the lethal yellow sample (red). (C-D) Uniform Manifold Approximation and Projection (UMAP) plots of the sequenced cells that passed quality control for each pairwise experiment. Among the cells isolated from nonagouti and lethal yellow paired samples (6,413 cells in C) and nonagouti and recessive yellow paired samples (4645 cells in D), skin cell types were identified and labeled as follows: Melanocytes (Mc), Dermal fibroblasts (DF), Dermal papilla (DP), Epidermis (Epi), Hair follicle and Outer root sheath (HF + ORS), and Transit amplifying and Matrix cells (TAC + Mx). Right panels show UMAP plots of the 2256 cells classified as non-cycling melanocytes. The nonagouti melanocytes (black) segregate from the recessive yellow and lethal yellow (gold) melanocytes consistent with a eumelanin and pheomelanin producing melanocytes having distinct transcriptional profiles.

Because the mitotic stage has a significant and influential effect on gene expression, all melanocytes outside the G_0_/G_1_ phase was removed using the same gene expression overlay approach with previously published cell cycle marker genes^21,22^ (Fig S3A, S3B). This defined the non-cycling melanocytes that were used for differential gene expression analysis comparison. Subsequent graph-based clustering analysis on the melanocyte population showed that both RHC mouse models displayed a bifurcation between the pheomelanin melanocytes and the eumelanin melanocytes captured in parallel, as displayed by UMAP (Fig 1C,D). Consistent with this, clustering of melanocytes by genotype was unique to melanocytes and not observed in the other cell types, suggesting that the effects of MC1R signaling loss are indeed restricted to melanocytes^23^.

Using this melanocyte population, differentially expressed genes (DEGs) (ANOVA; FDR < 0.05 and fold-change (FC) ±2) were identified in pair-wise experiments to compare lethal yellow to nonagouti melanocytes (Fig 1C) as well as recessive yellow to nonagouti melanocytes (Fig 1D). The differential lists from each experiment were intersected to identify shared genes which have consistent FC direction when comparing the nonagouti (*a/a*) melanocytes to the lethal yellow (*A^y^/a*) or recessive yellow (*Mc1r^e/e^*) melanocytes. These comparisons resulted in a set of 216 genes which we have identified as the Pheomelanin Gene Signature (PGS) (Table S2). We next asked to what extent these genes are associated with pigmentation by querying the PGS against a compiled list of genes annotated as causative for pigmentation phenotypes^24^ to assess the degree of overlap. Interestingly, only 5% of the PGS genes appeared in the previously annotated pigmentation gene list (Table 1).

**Table 1.** **Pigmentation phenotype associated genes found within the Pheomelanin Gene Signature (PGS)** with FDR and FC values for each pairwise experiment. In each pairwise experiment a negative FC indicates reduced expression in the pheomelanin producing melanocytes (lethal yellow, recessive yellow) compared to the eumelanin producing melanocytes isolated from nonagouti littermates.

| Gene | Lethal yellow |  | Recessive yellow |  |
| --- | --- | --- | --- | --- |
|  | FDR step up | Fold change | FDR step up | Fold change |
| *Tyrp1* | 0.00E+00 | -19.31 | 1.12E-67 | -83.90 |
| *Pmel* | 4.82E-46 | -2.33 | 5.82E-16 | -2.55 |
| *Mef2c* | 8.98E-40 | 2.69 | 1.93E-07 | 2.33 |
| *Oca2* | 1.77E-25 | -3.16 | 9.28E-05 | -2.47 |
| *Slc7a11* | 5.38E-23 | 7.72 | 3.79E-03 | 3.92 |
| *Irf4* | 1.32E-19 | -3.02 | 7.19E-03 | -2.61 |
| *Mc1r* | 2.58E-11 | 2.94 | 3.64E-03 | 3.63 |
| *Gas7* | 1.37E-07 | -2.28 | 1.05E-02 | -8.57 |
| *Sdc2* | 1.70E-07 | 2.39 | 8.94E-13 | 4.29 |
| *Pcbd1* | 1.25E-03 | 2.12 | 2.37E-02 | 2.34 |
| *Masp1* | 9.76E-03 | 3.11 | 1.73E-02 | 2.69 |

We next used PANTHER Gene Ontology (GO) and Ingenuity® Pathway Analysis (IPA) to identify significantly over-represented processes and pathways in the PGS. GO analysis revealed a prevalence of significant terms (FDR < 0.05) related to ‘morphogenesis’, ‘melanin biosynthesis’, and ‘neurogenesis’ / ‘neurotransmitter regulation’ (Fig 2A). Terms related to ‘melanin processes’ had equally high fold-enrichment; for example, 4 out of the 15 genes associated with ‘melanin biosynthetic process’ are present in the PGS (*Tyrp1*, *Mc1r*, *Oca2*, and *Pmel*). IPA highlighted canonical pathways related to ‘Neuroinflammation, ‘Synaptogenesis’ and ‘Neuron receptor signaling’ (Fig 2B). These terms are primarily driven by a host of receptors and ligands within the gamma-aminobutyric acid (GABA), N-methyl-D-aspartate (NMDA)/Glutamate, Ephrin, and Semaphorin families.

**Figure 2.**
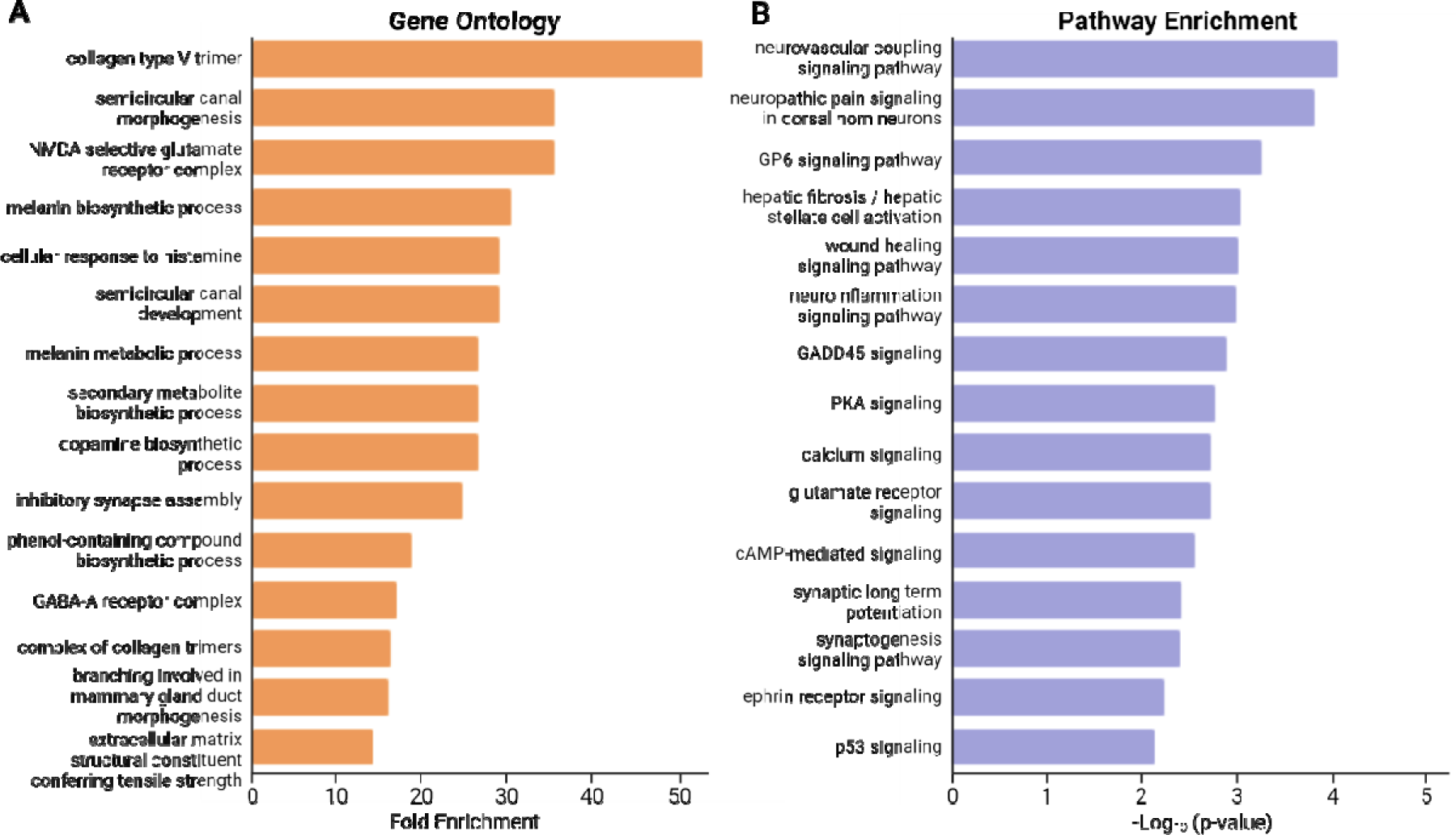
Enriched processes and pathways present in the PGS. (A) Top 15 significant GO terms ranked by fold-enrichment (FDR < 0.05). (B) Top 15 significant pathway terms from IPA ranked by significance (p-value <0.05).

### Predicted Transcriptional Regulators of the Pheomelanin Gene Signature include TBX3

To better identify the gene regulatory network regulating pheomelanin synthesis, we examined differentially expressed transcription factors, as well as predicted molecular upstream regulators of gene regulatory networks. The PGS contains 22 annotated transcription and co-transcription factors^25^ (Fig 3A). Many of these have established relationships to melanocyte and pigment cell biology, including neural crest development (*Hmga2*^26^, *Id4*^27^, *Mef2c*^28^*, Tbx3*^29^ and *Tfap2b*^30^), melanocyte differentiation (*Mef2c*^31^), melanogenesis (*Id4*^32^*, Irf4*^33^, and *Tbx3*^34,35^), oxidative stress protection (*Nr4a*^36^), and melanoma (*Hmga2*^37^*,Irf4*^38^, *Nfactc2*^39^, *Nr4a*^40^, *Tbx3*^41–45^ and *Zeb1*^46–48^). Of note, the *microphthalmia-associated transcription factor* (*Mitf*), a master melanocyte regulatory transcription factor^49^, was not identified as transcriptionally altered. Collectively, these data suggest that pheomelanin synthesis is not simply a result of eumelanin synthesis inactivation, but rather could involve multiple complex pathways downstream of MC1R signaling loss.

**Figure 3.**
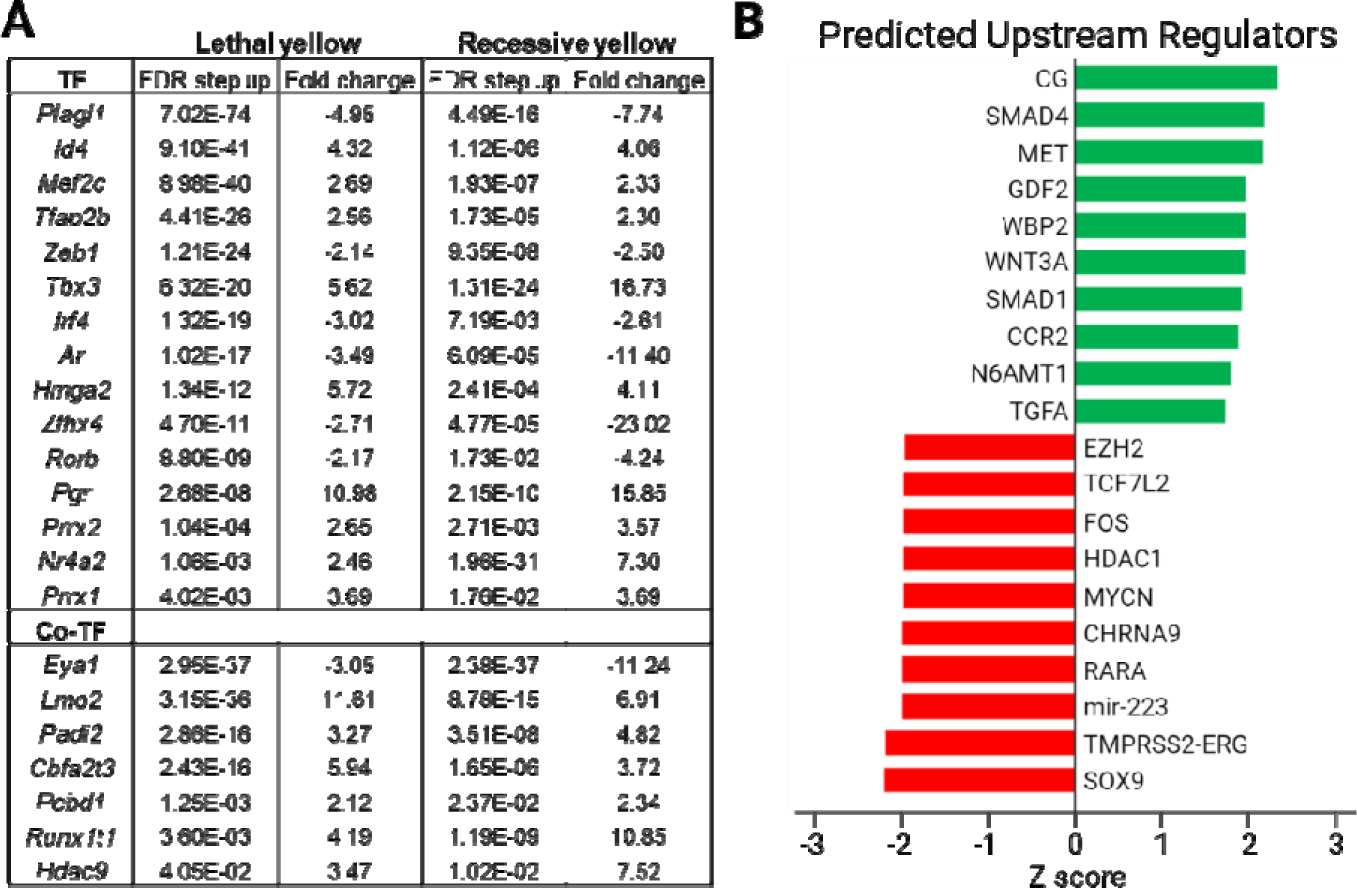
Identifying PGS regulatory factors. (A) List of all differentially expressed transcription/co-transcription factors found within the PGS and ranked by FDR from the nonagouti/lethal yellow pairwise differential analysis. (B) Top 10 predicted activated (green) and inhibited (red) regulators of the PGS gene list ranked by Z-score.

IPA was used to predict upstream regulators which may be driving expression of the PGS (Fig 3B). The top 10 predicted regulators include three molecules -- SMAD4, GDF2, and SMAD1 -- that are all related to TGFβ/BMP signaling. Importantly, TGFβ signaling can regulate melanin synthesis, with activation of the pathway resulting in hypopigmentation^50^. We reasoned that those transcriptional targets downstream of TGFβ/BMP signaling merited further analysis. Among these was *Tbx3*, a T-box family member activated by the TGF signaling pathway^51^. *Tbx3* showed β significantly increased expression in both the lethal yellow and recessive yellow samples (Fig 3A). TBX3 has previously been implicated in melanoma invasion^44^, but also has a key role as an anti-senescence transcription factor^1–4,45,52^. Up-regulation of TBX3 and consequent suppression of senescence in melanocytes lacking MC1R prior to acquisition of activating mutations in BRAF or NRAS would therefore provide a potential mechanism by which redheads would exhibit a predisposition to melanoma. In addition, TBX3 has previously been implicated in pigmentation, of Dun horses for example^53^, however, how TBX3 mediates these effects in unclear. Therefore, TBX3 was selected for further exploration with the goal of confirming TBX3 as a candidate transcription factor for direct regulation of key anti-senescence and melanogenesis target genes downstream from MC1R.

### TBX3 binds at T-box and E-box motifs

To assess the functional targets of TBX3 in melanocytes, Chromatin immunoprecipitation sequencing (ChIP-seq) was used to identify global targets of TBX3 binding. For this we used doxycycline-inducible TBX3 tagged with an HA epitope previously used to produce high-quality ChIP-seq datasets^54–56^ (Fig S4A). In our studies, the level of induction of TBX3 by doxycycline was adjusted to ensure physiological levels of TBX3-HA expression in the human melanoma cell line, 501mel (Fig S4B). We performed two replicate experiments and utilized MACS2 for peak calling (q-value cutoff ≤ 0.01). To identify common peaks between the two replicates, a filter was applied requiring an 80% overlap of coordinates between the 2 replicates, resulting in the identification of 20,449 common peaks (Fig S4C).

To determine the biological significance of these peaks, we used Genomic Regions

Enrichment Annotation Tool (GREAT)^57^ to identify 10,000 peaks associated with 7,109 candidate TBX3 gene targets (Methods; Table S3). We next assessed the binding distribution of these 10,000 peaks relative to gene transcriptional start sites (TSS). Promoter region binding within ±5kb of the TSS comprised only 12.68% of peaks, and the vast majority were situated ±20kb from the TSS, suggesting TBX3 binds distal enhancer elements (Fig 4A,B). GO enrichment for the 7,109 candidate TBX3 gene targets included multiple terms related to pigmentation/melanogenic pathways, as well as multiple terms associated with EMT, of which TBX3 is a well-known regulator (Fig 4C). Phenotype association analyses showed that pigmentation-related phenotypes were the most abundant for both human and mouse with a particular emphasis on hypopigmentation/abnormal melanocyte activity (Fig 4D).

**Figure 4.**
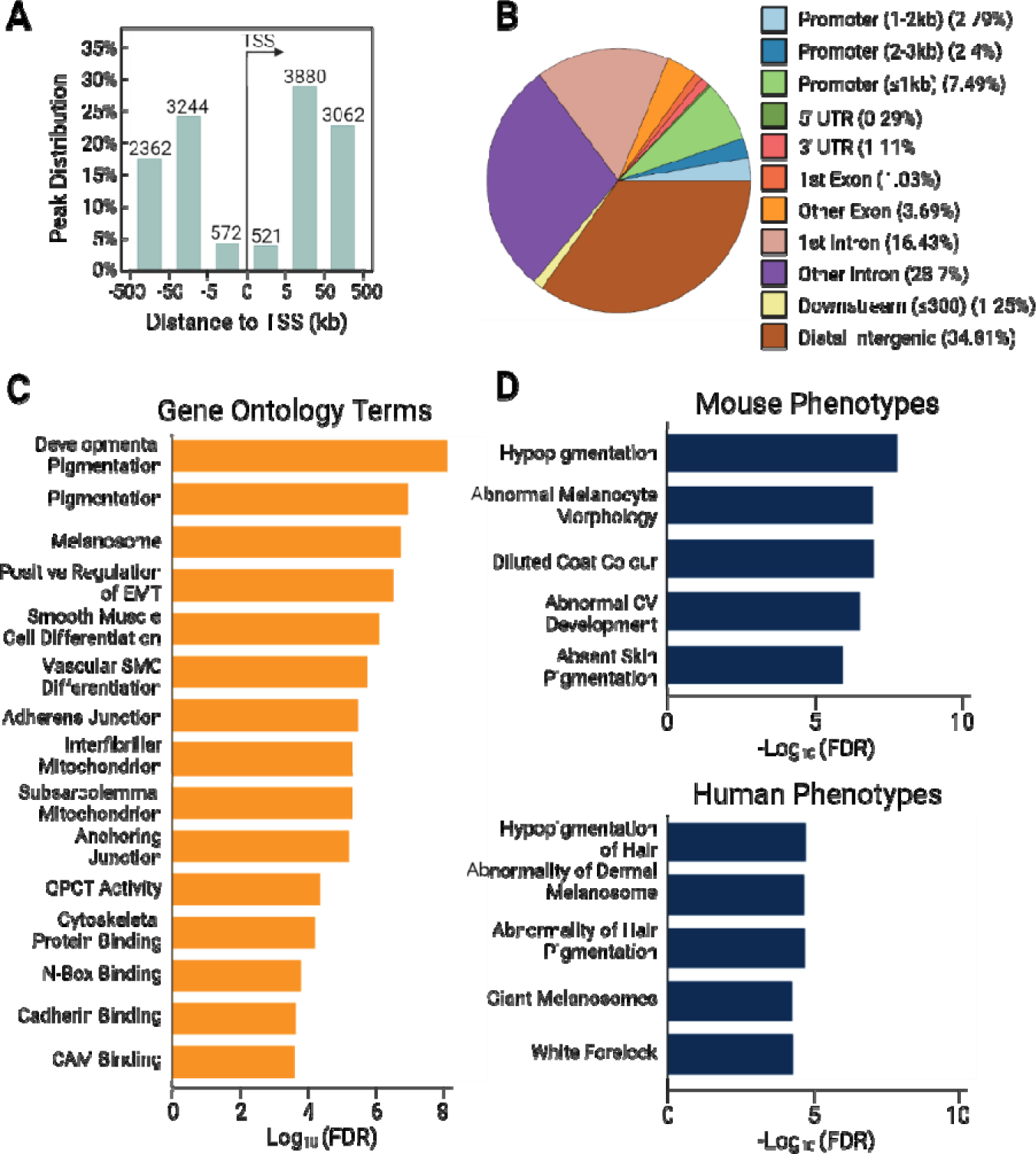
ChIP-seq analysis implicates TBX3 as a regulator of melanogenesis. (A) Position of called TBX3 peaks relative to the TSS. (B) ChIPseeker plot identifying genomic regions in which TBX3 is bound. (C) Top 15 GO terms of genes associated with TBX3 ChIP target genes ranked by −Log_10_ of the FDR. (D) Top 5 predicted phenotype associations (human and mouse) from TBX3 ChIP gene by −Log_10_ of the FDR.

We next used MEME-ChIP to assess motif enrichment within the TBX3 peaks. *De novo* motif analysis revealed that the top 10 sequences (JASPAR CORE vertebrates non-redundant v2), ranked by e-value, were not only associated with the expected T-box related elements, but also with E-box related motifs (Fig 5A). Interestingly, the palindromic canonical E-box sequence CACGTG is bound by basic-Helix-Loop-Helix (bHLH) and bHLHHzip transcription factors including MITF and its related family member. This sequence shares a 3bp GTG with the well-described core of the T-box element (AGGTGTGA) bound by T-box family members ^55–57^. Furthermore, TBX3 represses *CDH1* at a CAGGTGT site which shares 5bp with the canonical E-box^44^. Interestingly, a TBX3 peak does not appear at this site in our ChIP dataset; however, TBX3 and MITF share two peaks within intron 2, and another peak 1.6kb upstream of the promoter region (Fig S4D). Collectively, T-box and E-box motifs were found in the majority of TBX3-bound input sequences (8,320/10,000; 83.20%) with the E-box motifs occurring more frequently than the T-box motifs (69.17% vs 45.92%, respectively) (Fig 5B).

**Figure 5.**
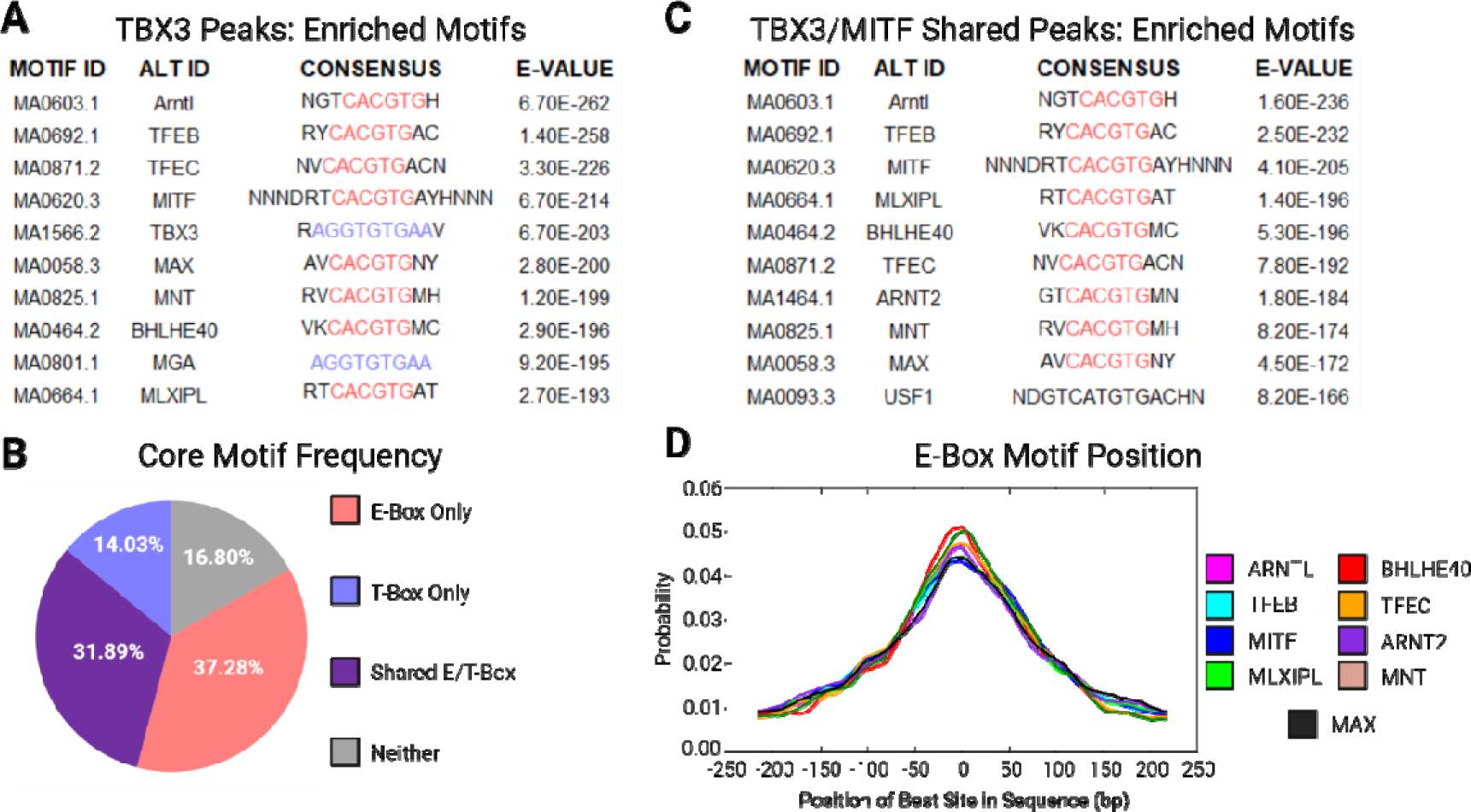
TBX3 binds MITF targets at E-boxes. (A) The top 10 motifs enriched within TBX3 ChIP peaks show over-representation of core E-box (red) and T-box (purple) sequences (B) The majority of TBX3 ChIP peaks contain either a T-box motif (14%), E-box motif (27%), or both (32%). (C) Top motifs of shared MITF and TBX3 peak sites (E-box = red; T-box = purple). (D) CentriMo plot showing top 9 centrally bound motifs.

Importantly, as the E-box motif is the primary recognition sequence associated with MITF^49^, we wanted to interrogate the degree of shared genome-wide binding sites between TBX3 and MITF. To assess for common occupancy of TBX3 and MITF we intersected our 20,449 TBX3 peaks with 12,137 called peaks from a previously published human MITF ChIP dataset in 501mel cells^58^. This yielded 4,758 common peaks which were assessed with MEME-ChIP for motif enrichment. The top 10 sequences (JASPAR CORE vertebrates non-redundant v2), ranked by e-value, were all E-box related motifs (Fig 5C) centrally located within the TBX3 bound peaks (Fig 5D). The frequency of TBX3 binding at centrally located E-box motifs also known to be occupied by MITF suggests that TBX3 and MITF may compete for binding at a set of shared regulatory elements.

To query the set of TBX3 and MITF dual candidate target genes, we utilized the same GREAT parameters as our TBX3 analysis to identify 7,095 MITF candidate target genes associated with peaks from the MITF ChIP dataset^59^ (Table S4). We then identified the mouse homologs for both the TBX3 and MITF target gene sets to allow comparison to the PGS gene list derived from *in vivo* mouse experiments (Table S5,S6). The intersection of TBX3 (6,499/7,109) and MITF (6,661/7,094) gene target lists identified a shared set of 4,072 genes, representing 62.6% of the TBX3 genome-wide target list (Fig 6A). Of these, 59 are found on the PGS (59/216; 27.3%), suggesting that a portion of the PGS is directly regulated by both TBX3 and MITF (Fig 6B/Table S7). Importantly, TBX3 and MITF shared binding sites at important pigmentation genes, with four of these having well-described MITF regulatory elements *IRF4, MC1R, OCA2,* and *TYRP1*^59–62^ (Fig 6C). Collectively, these data suggest that TBX3 can directly bind at E-box motifs associated with PGS genes, thus regulating pheomelanogenesis via modulation of MITF-mediated gene regulation.

**Figure 6.**
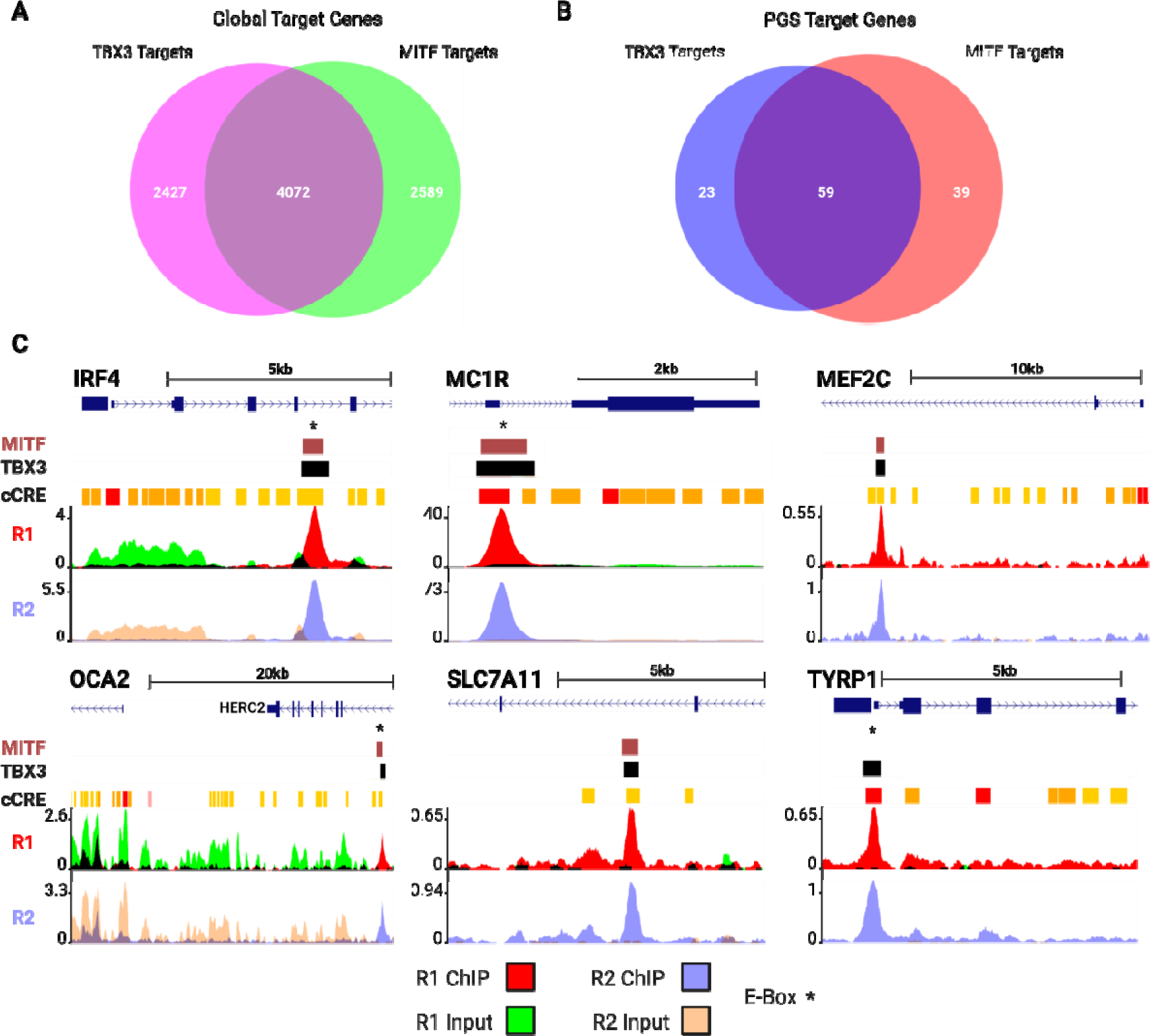
TBX3 binds the E-box of pigmentation gene targets. (A) Intersection showing the number of murine genes (4072) associated with ChIP peaks in both our TBX3 dataset and a published MITF dataset. (B) Intersection of PGS genes identified as targets of both TBX3 and MITF. (C) UCSC browser view of TBX3 and MITF ChIP peaks associated with pigmentation genes. Browser tracks are as follows: TBX3 = replicated binding sites in our dataset; MITF = ChIP peaks from Strub et al; cCRE=encode annotated candidate cis-regulatory elements. The location of verified MITF regulatory elements containing E-box motifs are indicated (*) (refs in text).

### TBX3 functions as a transcriptional activator and repressor

Our bioinformatic analyses strongly suggest that TBX3 can bind near PGS genes, including important genes related to melanogenesis. Traditionally, TBX3 has been shown to function as a transcriptional repressor; however, it is known to include an activator domain^52,63^. In MC1R-inhibited melanocytes, individual genes contained in the PGS exhibit either increased or decreased expression relative to eumelanin-producing nonagouti melanocytes. Therefore, we wanted to quantitatively analyze the extent to which TBX3 directly influences PGS expression levels. Bulk RNA-sequencing in human melanoma cells was performed following the transient reduction of TBX3, to determine (1) if the putative target genes were regulated by TBX3 and (2) if TBX3 repressed or activated these targets.

We performed transient knock-down of *TBX3* in 501mels for 24h, in triplicate, with two distinct siRNAs (TBX3_13.6 and TBX3_13.8) and a non-coding control. At 24 hours we collected lysates, harvested RNA, and then submitted samples for bulk RNA-seq. We then performed differential analysis and identified significant DEGs (padj < 0.05) that were responsive to TBX3 depletion (Table S8). We next intersected DEGs from the two small interfering RNAs and removed genes in which FC occurred in contradictory directions, resulting in 1,420 positive FC genes (+FC) and 1,340 negative FC genes (-FC) (Fig 7A). To identify candidate genes regulated by TBX3, we intersected this list with our TBX3 ChIP target genes, which identified a total of 1,091 targets (623 +FC, 468-FC) that show both responsiveness to TBX3 depletion as well as TBX3 binding near their respective genomic loci (Table S9).

**Figure 7.**
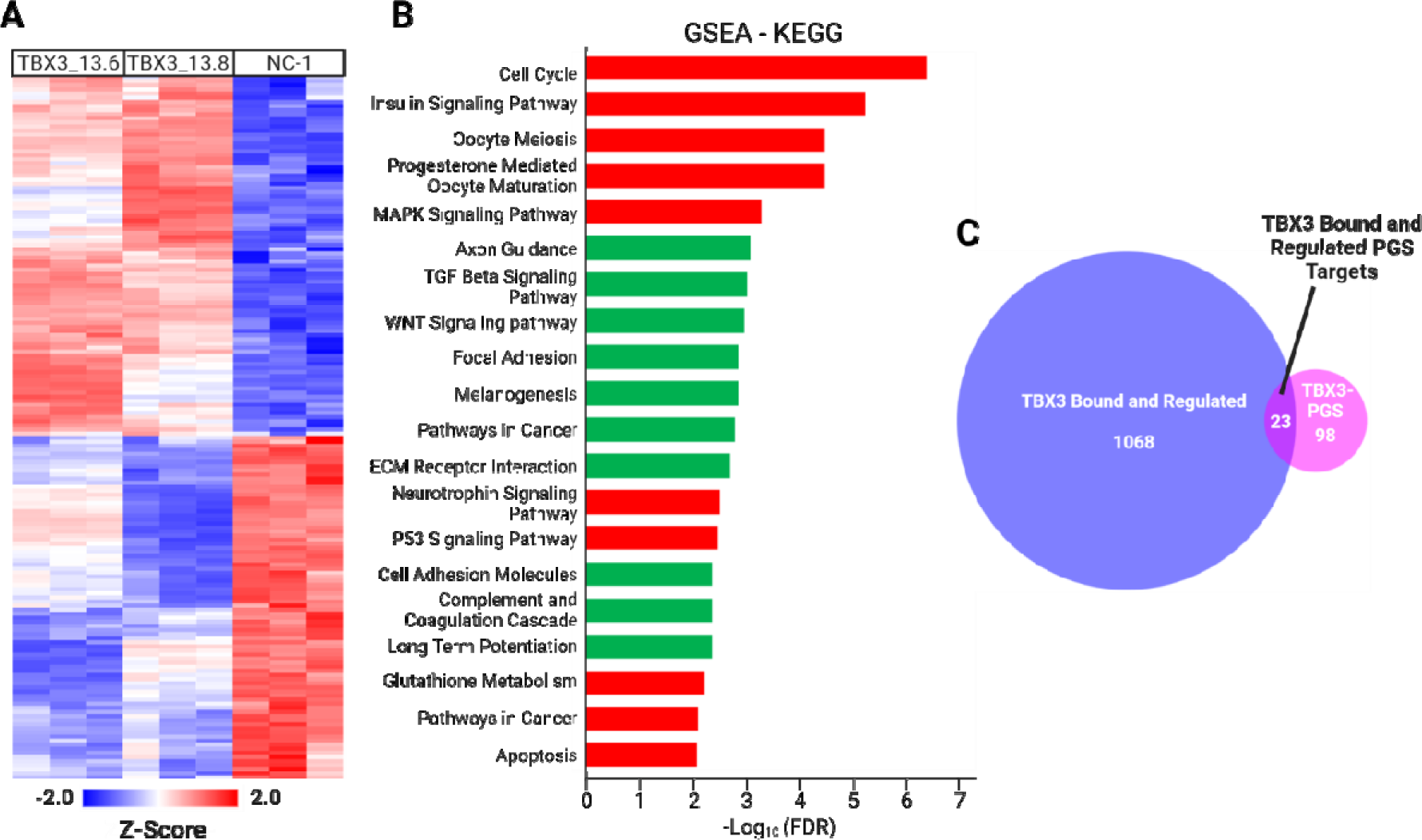
TBX3 binds to and regulates PGS genes. (A) Heatmap depicting relative gene expression of 2780 DEG (Adjusted p-value < 0.05) from the TBX3 knockdown samples against a non-coding control. (B) Top KEGG terms ranked by −Log_10_ of the p-value for upregulated (green) and downregulated (red) genes. (C) Intersection of DE TBX3 gene targets against PGS TBX3 targets.

### TBX3 regulates PGS and cell cycle genes

We next performed Gene Set Enrichment Analysis (GSEA) on the candidate direct TBX3 targets to identify significant (FDR < 0.05) KEGG terms associated with the +FC and -FC genes. Terms for the +FC list include those related to TBX3’s known role in melanoma, such as ‘TGFb signaling pathway’, ‘focal adhesion’, and ‘ECM receptor interaction’ (Fig 7B). Importantly, ‘melanogenesis’ was one of the most significant terms. Enriched terms for -FC genes include those consistent with TBX3’s pro-proliferative and anti-senescent functions, such as ‘cell cycle’, MAPK signaling pathway’, and ‘p53 signaling pathway’ (Fig 7B). Specifically, this includes cell cycle progression genes *CDK1* and *CDK2* (Table S9). Lastly, we wanted to assess which PGS targets were specifically bound and regulated in our *in vitro* system. We first converted the 1,091 +/-FC genes bound and regulated by TBX3 to their known mouse homologs, yielding 1,041 genes. These were intersected with the 98 genes comprising TBX3 targets of the PGS, of which 23 were present (Fig 7C). Collectively, these data suggest that TBX3 can regulate melanogenesis as both a transcriptional activator and repressor of PGS genes.

## Discussion

Coding variants in *MC1R* are associated with the RHC trait. However, the gene regulatory networks (GRNs) downstream of MC1R remain poorly understood. Through a high-resolution, single-cell RNA-sequencing approach we have identified a Pheomelanin Gene Signature in melanocytes comprising candidate genes which may regulate both pheomelanogenesis and melanomagenesis. One such candidate is *Tbx3* which we show can bind and regulate PGS genes. Interestingly, we have shown that the transcriptional regulatory activities of TBX3 include activation and repression of candidate genes. Furthermore, this function is not restricted to DNA binding via the T-box motif, rather it includes binding the canonical CACGTG E-box, which is the main recognition sequence of the melanocyte master regulatory transcription factor MITF.

The vast majority of genes comprising the PGS have no previously established ties to pigmentation phenotypes (205/216; 94.9%). A common approach to begin investigation of a novel gene list involves identifying over-enriched gene families and pathways. One such PGS-enriched family encompasses neurogenesis and neurotransmitter regulation and is primarily driven by receptors and ligands within the gamma-aminobutyric acid (GABA), N-methyl-D-aspartate (NMDA)/Glutamate, Ephrin, and Semaphorin families. Mature melanocytes show morphological similarities with neurons in that they are highly dendritic, which enables contact with multiple keratinocytes for melanin transfer. Furthermore, normal human melanocytes have GO enrichment for ‘neuron ensheathment and synapse’^22^. Interestingly, melanocytes that predominantly produce pheomelanin in the presence of ASIP lack dendricity^64^. Likewise, inhibition of glutamate signaling through NDMA receptors has been shown to inhibit melanocyte filipodia attachment to keratinocytes as well as reduce melanosome transfer efficacy^65^. A recent study has shown that GABA signaling between keratinocytes and melanoma cells occurs through neuronal like synapses, and further shows that this communication promotes melanoma initation^66^. Interestingly, in this context, keratinocytes showed increased GABA receptor expression while melanoma cells did not. Collectively, increased expression of these gene families may reflect both intrinsic and extrinsic signaling to promote melanocyte development, dendrite formation, and melanosome transfer abilities.

Previous work has shown that the benzodiazepine Diazepam, which works to increase the action of GABA, increases both melanogenesis and melanocyte dendricity^67^. Likewise, NMDA receptors appear to promote melanin synthesis and tyrosinase activity in the cuttlefish, *Sepia officinalis*^67,68^. In addition, glutamate receptor genes, especially *Grm1*, are expressed in human melanomas^69^, and ectopic expression of *mGluR1* in melanocytes leads to oncogenic transformation and metastatic spread with 100% penetrance^69^. Ephrins play an essential role in regulating biological processes such as cell-cell attachment, cell shape, and motility^70^. Along with Semaphorins, they are expressed in neural crest cells (which give rise to melanocytes) and are of vital importance in regulating neural crest migration^71^. Genes within these families also show differential expression in melanocytes treated with ASIP, suggesting that this treatment results in dedifferentiation towards a neural crest-like state^72^. Our data further validate this idea and suggest that MC1R acts downstream of ASIP to effect these changes, as loss of MC1R in primary melanocytes drives similar gene expression changes.

Transcription factors play an essential role in the regulation of gene expression. The most well-characterized of these in melanocytes is *Mitf.* This gene encodes a master regulatory transcription factor implicit in melanocyte specificity, proliferation, survival, migration, differentiation, and melanogenesis, and whose expression has been reported to lie downstream from MC1R signaling^73,74^. *Mitf* does not exhibit differential expression in our study, nor do many of its known post-transcriptional/translational modifiers, such as MAPKs, *Gsk3b*, *p300*, and *Stat3,* although changes in the activity of these signaling molecules may not be apparent in the gene expression profile. Similarly, other well-characterized transcription factors in melanocyte biology, including *Sox10*, *Pax3*, *Lef1*, and *Creb,* which can all regulate *Mitf,* also exhibit no differential expression. This supports that the primary transcriptional changes driving activation of pheomelanin, and repression of eumelanin, synthesis downstream of MC1R signaling are not mediated through reduced *Mitf* transcription, but instead through transcriptional regulation of other factors that modulate the DNA binding or post-transcriptional regulation of MITF.

One of the primary candidate genes we performed further analysis on is the T-box transcription factor family member, *Tbx3*. This candidate stood out for the following reasons: (1) it is activated by TGFβ/BMP signaling^51,63,75^, (2) it has high fold-enrichment in melanocytes from both pheomelanin-producing mouse genotypes (Fig 3A), (3) it is found across multiple terms from our GO analysis, (4) it has known functional roles in developmental states of melanocytes^29,34,35^ (5) it regulates developmental processes across multiple tissue/cell types, including heart^76–78^, lungs^79^, limbs^80,81^, inner ear^82^, and neural crest^29,76^, and (6) is highly expressed in a multitude of cancers, especially melanoma, where its primary roles appear to be in metastasis by activation of EMT, at least in part through down-regulation of E-cadherin, as well as suppression of senescence^41,43–45,83,84^.

The differential expression of *Tbx3* in eumelanin- and pheomelanin-expressing melanocytes, along with genome-wide binding of TBX3 to T- and E-box elements in melanoma cells, suggested that TBX3 may directly regulate MITF targets downstream of MC1R signaling. Through ChIP and bulk RNA-seq we’ve shown that TBX3 can bind and regulate genes within the PGS. Although TBX3 is often viewed as a transcriptional repressor, it has been shown to activate gene expression of putative target genes^52^. Likewise, our data strongly suggests that TBX3 can both activate or repress target genes. These targets include genes with known associations to pigmentation phenotypes. An intriguing example of TBX3-mediated activation is *Slc7a11*. This gene encodes the cystine/glutamate exchanger (xCT) which imports cystine into the cell where it is reduced into two molecules of cysteine, a necessary component for pheomelanin synthesis^85,86^. The increased pool of cysteine provides an environment that is favorable for pheomelanin synthesis. Increased expression of *Slc7a11* by TBX3 suggests an active process downstream of MC1R to favor pheomelanin synthesis, distinct from the repression of eumelanin synthesis. The importance of cysteine goes beyond pheomelanin synthesis, as it is an essential factor of the glutathione system and oxidative stress regulation as well as ferroptosis inhibiton^86,87^. Following *TBX3* knockdown we observed a significant reduction of *SLC7A11* expression, with associated enrichment of pathways associated with ‘glutathione metabolism’ and ‘apoptosis’. Given the cytotoxicity of pheomelanin, regulation of cysteine levels within melanocytes is critical and presents a paradox where cysteine is essential for regulating oxidative stress; however, too much can exacerbate cell stress through pheomelanin accumulation.

TBX3 is implicated in senescence by-pass partly through its ability to repress cell cycle checkpoint genes such as CDKN2B^52^. Our *TBX3* knockdown data support this, with *CDKN2B* showing significantly increased expression. Furthermore, our data also support a role for TBX3 in senescence by-pass through activation of cell cycle progression genes, where the loss of *TBX3* yields decreased expression of *CDK1* and *CDK2* (Table S9). Willmer et al^88^ have shown that TBX3 is important in S-phase transition in part due to post-translational phosphorylation via CDK2^88^. More recent work from this group suggests a more complex relationship, whereby TBX3 itself promotes *CDK2* expression^89^. Our data further support this model and suggest a further way in which TBX3 can contribute to oncogenic transformation by facilitating senescence bypass. These observations may pave the way in explaining why redheads with diminished MC1R function may be predisposed to melanoma.

The T-box family members comprise five sub-groups, with TBX3 as a member of the TBX2 sub-group. TBX2 and TBX3 share 95% homology within the DNA binding domain^63^. TBX2, which canonically binds the T-box, can also bind the E-box motif, and functions as both an activator and repressor^56^. Our data support the idea that TBX3 also binds to, and regulates gene expression, via the E-box motif. However, unlike TBX2, the type of motif bound does not seem to influence its action (Fig S5A). E-box site recognition of TBX3 can possibly be explained with protein structural studies. As mentioned previously, the canonical E-box and T-box elements share a 3bp (GTG) region. This corresponds to base pairs 3-5 of the T-box (AGGTGTGA). Consequently, this 3bp core significantly affects TBX3’s ability to bind DNA, whereas the flanking base pairs are deemed less crucial^90^. As a monomer, TBX3 can also bind to either the major or minor groove^90^. Therefore, while bound in the minor grove, it is possible that TBX3 may interact with other transcription factors bound in the major grove, including MITF. The degree to which TBX3 modulates gene expression is most likely dependent on several factors, including, cell type, differentiation state, co-factor availability, chromatin accessibility, and biological state (e.g., methylated, acetylated). Variation across these factors could explain why more putative genes targets regulated by TBX3 in human melanoma cells *in vitro* are not represented in the *in vivo* healthy murine melanocyte PGS.

In this study, we aimed to profile the transcriptome in melanocytes with aberrant MC1R signaling. To this end we have defined a set of genes (PGS) that merit further investigation towards better understanding the complex GRNs downstream of MC1R signaling. This includes the melanoma-associated transcription factor, *Tbx3*. Furthermore, we believe the Pheomelanin Gene Signature provides important candidate genes that merit further follow-up regarding the regulation of melanocyte development, melanogenesis, and melanomagenesis.

## Materials and Methods

### Single-Cell RNA-Sequencing and Differential Expression Analysis

FACS-isolated melanocytes were captured in a solution of 1xPBS + 0.04% FBS. Cell viability and concentration were analyzed using a Luna-FX7 cell counter (Logos Biosystems). Gel Bead-in-Emulsions were prepared by loading 5,000 cells/sample onto a Chromium Chip G (10x Genomics #1000127) and run using the Chromium controller (10x Genomics #1000204). cDNA libraries were generated with Chromium Single Cell 3’ GEM, Library and Gel Bead Kit V3.1 (10x Genomics #1000128). Samples were indexed and pairs for each experiment were pooled for sequencing using the NextSeq 500/550 Hi Output KT v2.5 (Illumina #20024907) on an Illumina NextSeq550. Barcoded reads were aligned to the mm39 genome using STAR aligner. Downstream processing of the data was performed with Partek Flow™ Genomic Analysis Software.

Single cell QA/QC included steps to filter out cells which did not meet the following parameters defined in the two pairwise experiments (lethal yellow / recessive yellow) as: barcode count 600-32,000/3,000-45,000, number of genes detected 500-6000/500-7000, and % ribosomal counts 12-43%/12-39%. Of the 31,017 annotated murine genes, 16,793 (54.14%)/20,060 (64.7%) passed an exclusion cutoff where genes were excluded if they had zero counts in at least 99.9% of the cells. The remaining read counts were normalized using a log_2_-based method on Counts Per Million (CPM). Normalized counts underwent dimensional reduction via a Principal Components Analysis (PCA) on the first 100 principal components and unsupervised, non-biased, Graph-based clustering through a Louvain algorithm on the first 20 principal components. Graph-based clusters underwent further dimensional reduction for visualization on a Uniform Manifold Approximation and Projection diagram. Postnatal mouse skin cell types including dermal fibroblasts, dermal papilla, transit amplifying cells, matrix cells, and epidermal cells were identified by utilizing published signature gene lists^20^. The most highly expressed genes (top 10%) from the signature gene lists were overlayed on the UMAP and the graph-based clusters which coincided with expression overlay were assigned to each cell type. All non-melanocyte cell types were subsequently filtered out, leaving only clusters classified as melanocytes for downstream analysis. The melanocyte only population underwent a further iteration of PCA, graph-based clustering, and UMAP display. To eliminate the variable of cellular heterogeneity as influenced by the mitotic state, we selected for cells within the G_0_ phase using a gene expression overlay established by Hsiao et al^21^. The melanocytes within the G_0_ phase underwent a further iteration of PCA, graph-based clustering, and UMAP analysis. Differential gene expression lists of melanocytes were generated using the Partek ANOVA differential expression analysis. Genes considered statistically significant had a False-discovery rate of <0.05 and a fold-change of ±2.

### Gene ontology and pathway analysis

Genes comprising the Pheomelanin Gene Signature were uploaded to the Gene Ontology Resource for GO Enrichment Analysis via PANTHER for Overrepresentation Tests (GO Ontology database DOI:10.5281/zenodo.6399963; Released 2022-03-22). The PGS was run against a reference list comprising all *Mus musculus* genes in the database. Fisher’s Exact test was performed with a False Discovery Rate correction for GO Biological Process Complete, GO Cell Component Complete, and GO Molecular Function Complete with significant results being those with an FDR < 0.05. To identify predicted upstream regulators, we utilized QIAGEN’s Ingenuity® Pathway Analysis (73620684 (Release Date: 2022-03-12). Testing was done via Expression Core Analysis on the observed experimental fold change of the PGS for both pairwise experiments.

### Gene Set Enrichment Analysis

Gene lists comprising the significant DE +FC and -FC genes regulated by TBX3 were uploaded to the Molecular Signatures Database (MSigDB) for gene set enrichment analysis utilizing the Kyoto Encyclopedia of Genes and Genomes (KEGG). For the +FC list, the top 500 genes by fold-change for TBX3_13.8 were used as the maximum input value is 500 genes. Each list was tested independently with significant terms being those with an FDR q-value < 0.05.

### TBX3 ChIP Analysis

Prior to mapping to the human reference genome (hg38) with Bowtie2 (v.2.3.5)^91^, quality of the raw sequencing data was evaluated using FastQC (v.0.11.7), with adapter contamination removed using CutAdapt (v.2.8). Peak calling was performed using MACS2 (v.2.1.2)^92^ with a 0.01 q-value threshold. Bowtie2-generated SAM files were compressed to BAM files, indexed using SAMtools (v.1.9), and sequentially converted to bigWig files using USCStools (v.373). The BED files of both replicates were intersected using the UCSC Table Browser to pull out common peaks from replicate 2 (40,974) that had ≥80% overlap with replicate 1 (32,479), which yielded a group of 20,449 common peaks. For simplicity of downstream analysis, the 20,449 peak coordinates used were derived from the BED position parameters of replicate 1.

The Genomic Regions Enrichment of Annotations Tool (GREAT v4.0.4)^57^ was used to analyze the functional significance of cis-regulatory regions of the TBX3 bound ChIP peaks. Each annotated gene was assigned a basal regulatory domain of 5kb upstream, and 1kb downstream, of the transcriptional start site. The regulatory domain of that gene was then extended in both directions to the nearest gene’s basal domain but with no more than a 100kb extension in one direction^57^. Discovery of enriched binding motifs was done using the MEME-ChIP^93^ DNA binding motif analysis tool. 250bps were added up and downstream of the chromosomal coordinates corresponding to the absolute summit of all overlapping peaks. A FASTA file was generated using the usegalaxy.org bedtools application ‘GetFastaBed’^94^. This was run against the default settings in MEME-ChIP, except for maximum width for ‘What width motifs should MEME and STREME find?’ which was increased to 20 bases. Peak distributions were obtained from GREAT and ChIPseeker^95^, via usegalaxy.org. The histogram was produced by GREAT, while the venn diagram was created with ChIPseeker.

### Bulk RNAseq of *in vitro* 501mel Human Melanoma Cells

501mel cells (140,000/well) were plated in a 12-well plate (1.2mL/well). dsiRNAs (NC1, TBX3 13.6, TBX 13.8; Integrated DNA Technologies) were diluted in 300uL of Opti-MEM (final concertation of 20nM). Lipofectamine RNAiMAX (6uL/100uL; Invitrogen 13778075) was then diluted in 300uL of Opti-MEM. Diluted dsiRNA and Lipofectamine were mixed and set to incubate at RT for 5 mins. 200ul of dsiRNA-Lipofectamine mix was added to each well. At 24h, cells were lysed for RNA collection via (Zymo Research R2053). RNA purity was assessed via tapestation with all samples having a RIN purity of ≥ 9.0. RNA samples were submitted to Omega Bioservices (Norcross, GA, USA) for bulk RNAseq. RNA purity was again assessed via an Agilent tapestation with all samples having a RIN ≥ 9.0. Library construction was completed using NEBNext® Ultra™ II Directional RNA Library Prep Kit for Illumina®. Samples were sequenced using NovaSeq 6000 (Illumina) PE150 with a ∼40M (20M in each direction) read throughput for each sample. Phred Mean Quality scores for all samples was ≥ 38.86. Data were analyzed by ROSALIND® (https://rosalind.bio/), with a HyperScale architecture developed by ROSALIND, Inc. (San Diego, CA).

### Figure Construction

Figures were assembled and created with BioRender.com. UMAPs were generated with Partek Flow. Venn diagrams were produced using DeepVenn. Binding distribution, Fig 3A, was modified from original GREAT output, while Fig 3B was modified from the ChIP seeker output from usegalaxy.org. Motif distribution for Fig 5D is modified from the MEME CentriMo tool.

## Author Contributions

Contribution: W.J.P., D.E.W.-C, H.M.B, P.M.A., and C.R.G. designed the research study; H.M.B., D.E.W.-C, S.L., P.L., K.M.B., T.Z. acquired the data; H.M.B., D.E.W.-C, K.M.B., T.Z. analyzed data; H.M.B, D.E.W.-C, C.R.G., and W.J.P. wrote the manuscript; H.M.B., D.E.W.-C., S.L., P.L., C.R.G., P.M.A., and W.J.P. edited the manuscript; and all authors approved the final manuscript.

## Data Availability

Original data created for the study are or will be available in a persistent repository upon publication. TBX3 ChIP is deposited in Gene Expression Omnibus (GEO) as GSE223256; Currently compiling pertinent information for submission of bulk and single-cell RNA-seq datasets to GEO.

## Acknowledgments

We are grateful to Dr. Jonathan Zippin for providing the C57BL/6J-Mc1r^e^/J mice and helpful scientific discussion. We further extend thanks to the NHGRI Microarray (Dr. Abdel G. Elkahloun and Bayu Sisay) and Flow Cytometry (Stacie Anderson and Martha Kirby) cores. Lastly, we greatly appreciate the help of Dr. Laura Baxter in providing helpful editing and feedback.

## Supporting Information

**S1 Fig. Classification Scheme for nonagouti and lethal yellow cells.** (A) Uniform Manifold Approximation and Projection (UMAP) plot of lethal yellow and nonagouti cells classified by skin cell type: Melanocytes (Mc), Dermal fibroblasts (DF), Dermal papilla (DP), Epidermis (Epi), Hair follicle and Outer root sheath (HF + ORS), and Transit amplifying and Matrix cells (TAC + Mx). (B) UMAP of cells classified by graph-based cluster. (C) UMAPs depicting heatmap of cell type scoring.

**S2 Fig. Classification Scheme for nonagouti and recessive yellow cells.** (A) Uniform Manifold Approximation and Projection (UMAP) plot of the recessive yellow and nonagouti cells classified by skin cell type: Melanocytes (Mc), Dermal fibroblasts (DF), Dermal papilla (DP), Epidermis (Epi), Hair follicle and Outer root sheath (HF + ORS), and Transit amplifying and Matrix cells (TAC + Mx). (B) UMAP of cells classified by graph-based cluster. (C) UMAPs depicting heatmap of cell type scoring.

**S3 Fig. Melanocytes cell cycle.** (A) Total melanocytes from lethal yellow and nonagouti pairwise experiment (left), cells within the cell cycle highlighted in blue (right). (B) Total melanocytes from recessive yellow and nonagouti pairwise experiment (left), cells within the cell cycle highlighted in blue (right).

**S4 Fig. Inducible TBX3 cell line validation.** (A) schematic of piggyBAC transposase system with Tet-on inducible expression of TBX3. Dox, doxycycline; TRE, tetracycline response element; rtTA, reverse tetracycline-controlled transactivator. (B) Western blot of polyclonal 501mel-iTBX3-3xHA-PuroR inducible cells treated with varying amounts of doxycycline for 24h. (C) Venn diagram depicting shared peaks between both replicate ChIP experiments. (D) UCSC browser view of TBX3 and MITF ChIP peaks associated with E-cadherin. Browser tracks are as follows: TBX3 = replicated binding sites in our dataset; MITF = ChIP peaks from Strub et al; cCRE=encode annotated candidate cis-regulatory elements.

**S5 Fig. Motif enrichment by fold-change direction in TBX3 KD studies.** (A) Top 10 motifs, ranked by e-value, of TBX3 ChIP peaks associated with putative target genes with increased expression (left) or decreased expression (right) upon TBX3 knock-down.

**S1-S5 Figure File. Berns et al_Supplementary Figures S1-S5.doc.**

**S1 Table. Gene Signature of Murine Skin Cells from Rezza et al.**

**S2 Table. Pheomelanin Gene Signature.**

**S3 Table. GREAT Output for TBX3 ChIP Dataset.**

**S4 Table. GREAT Output for MITF ChIP Dataset.**

**S5 Table. Human to Mouse Conversion of TBX3 Target Genes.**

**S6 Table. Human to Mouse Conversion of MITF Target Genes.**

**S7 Table. Gene Target Overlap.**

**S8 Table. Significant Differentially Expressed Genes from TBX3 Knockdown.**

**S9 Table. Significant Differentially Expressed Genes from TBX3 Knockdown which are TBX3 Targets Identified by ChIP.**

**S1-S9 Tables File. Berns et al_**Supplementary Tables S1-S9**.xls.**

## References

1.) Ha, L., Merlino, G. & Sviderskaya, E. V. Melanomagenesis: Overcoming the barrier of melanocyte senescence. Cell Cycle 7, 1944–1948 (2008).

2.) Bennett, D. C. & Medrano, E. E. Molecular Regulation of Melanocyte Senescence. Pigm Cell Res 15, 242–250 (2002).

3.) Vredeveld, L. C. W., et al. Abrogation of BRAFV600E-induced senescence by PI3K pathway activation contributes to melanomagenesis. Gene Dev 26, 1055–1069 (2012).

4.) Wagstaff, W., et al. Melanoma --- Molecular genetics, metastasis, targeted therapies, immunotherapies, and therapeutic resistance. Genes Dis 9, 1608–1623 (2022).

5.) Rhodes, A. R., Weinstock, M. A., Fitzpatrick, T. B., Mihm, M. C. & Sober, A. J. Risk Factors for Cutaneous Melanoma: A Practical Method of Recognizing Predisposed Individuals. Jama 258, 3146–3154 (1987).

6.) Rayner, J. E., et al. Phenotypic and genotypic analysis of amelanotic and hypomelanotic melanoma patients. J Eur Acad Dermatol 33, 1076–1083 (2019).

7.) Schiöth, H. B., et al. Loss of Function Mutations of the Human Melanocortin 1 Receptor Are Common and Are Associated with Red Hair. Biochem Bioph Res Co 260, 488–491 (1999).

8.) Pavan, W. J. & Sturm, R. A. The Genetics of Human Skin and Hair Pigmentation. Annu Rev Genom Hum G 20, 1–32 (2019).

9.) Mort, R. L., Jackson, I. J. & Patton, E. E. The melanocyte lineage in development and disease. Development 142, 620–632 (2015).

10.) Napolitano, A., Panzella, L., Monfrecola, G. & d’Ischia, M. Pheomelanin-induced oxidative stress: bright and dark chemistry bridging red hair phenotype and melanoma. Pigm Cell Melanoma R 27, 721–733 (2014).

11.) Mitra, D., et al. An ultraviolet-radiation-independent pathway to melanoma carcinogenesis in the red hair/fair skin background. Nature 491, 449–453 (2012).

12.) Robles-Espinoza, C. D., et al. Germline MC1R status influences somatic mutation burden in melanoma. Nat Commun 7, 12064 (2016).

13.) Cao, J., et al. MC1R Is a Potent Regulator of PTEN after UV Exposure in Melanocytes. Mol Cell 51, 409–422 (2013).

14.) HAUSCHKA, T. S., JACOBS, B. B. & HOLDRIDGE, B. A. Recessive Yellow and its Interaction With Belted in the Mouse. J Hered 59, 339–341 (1968).

15.) Michaud, E. J., Bultman, S. J., Stubbs, L. J. & Woychik, R. P. The embryonic lethality of homozygous lethal yellow mice (Ay/Ay) is associated with the disruption of a novel RNA-binding protein. Gene Dev 7, 1203–1213 (1993).

16.) Michaud, E. J., et al. A molecular model for the genetic and phenotypic characteristics of the mouse lethal yellow (Ay) mutation. Proc National Acad Sci 91, 2562–2566 (1994).

17.) April, C. S. & Barsh, G. S. Skin layer-specific transcriptional profiles in normal and recessive yellow (Mc1re/Mc1re) mice. Pigm Cell Res 19, 194–205 (2006).

18.) Haltaufderhyde, K. D. & Oancea, E. Genome-wide transcriptome analysis of human epidermal melanocytes. Genomics 104, 482–489 (2014).

19.) Harris, M. L., et al. A direct link between MITF, innate immunity, and hair graying. Plos Biol 16, e2003648 (2018).

20.) Rezza, A., et al. Signaling Networks among Stem Cell Precursors, Transit-Amplifying Progenitors, and their Niche in Developing Hair Follicles. Cell Reports 14, 3001–3018 (2016).

21.) Hsiao, C. J., et al. Characterizing and inferring quantitative cell cycle phase in single-cell RNA-seq data analysis. Genome Res 30, 611–621 (2020).

22.) Belote, R. L. et al. Human melanocyte development and melanoma dedifferentiation at single-cell resolution. Nat Cell Biol 1–13 (2021) doi:10.1038/s41556-021-00740-8.

23.) Guida, S., Guida, G. & Goding, C. R. MC1R Functions, Expression, and Implications for Targeted Therapy. J Invest Dermatol (2021) doi:10.1016/j.jid.2021.06.018.

24.) Baxter, L. L., Watkins-Chow, D. E., Pavan, W. J. & Loftus, S. K. A curated gene list for expanding the horizons of pigmentation biology. Pigm Cell Melanoma R 32, 348–358 (2019).

25.) Hu, H., et al. AnimalTFDB 3.0: a comprehensive resource for annotation and prediction of animal transcription factors. Nucleic Acids Res 47, gky822 (2018).

26.) Macrì, S., et al. Hmga2 is required for neural crest cell specification in Xenopus laevis. Dev Biol 411, 25–37 (2016).

27.) Kee, Y. & Bronner-Fraser, M. Id4 expression and its relationship to other Id genes during avian embryonic development. Mech Develop 109, 341–345 (2001).

28.) Hu, J., et al. Endothelin signaling activates Mef2c expression in the neural crest through a MEF2C-dependent positive-feedback transcriptional pathway. Development 142, 2775– 2780 (2015).

29.) López, S. H., et al. Loss of Tbx3 in murine neural crest reduces enteric glia and causes cleft palate, but does not influence heart development or bowel transit. Dev Biol 444, S337–S351 (2018).

30.) Seberg, H. E., Otterloo, E. V. & Cornell, R. A. Beyond MITF: Multiple transcription factors directly regulate the cellular phenotype in melanocytes and melanoma. Pigm Cell Melanoma R 30, 454–466 (2017).

31.) Agarwal, P., et al. The MADS box transcription factor MEF2C regulates melanocyte development and is a direct transcriptional target and partner of SOX10. Development 138, 2555–2565 (2011).

32.) DiVito, K. A., Trabosh, V. A., Chen, Y., Simbulan-Rosenthal, C. M. & Rosenthal, D. S. Inhibitor of differentiation-4 (Id4) stimulates pigmentation in melanoma leading to histiocyte infiltration. Exp Dermatol 24, 101–107 (2015).

33.) Chhabra, Y., et al. Genetic variation in IRF4 expression modulates growth characteristics, tyrosinase expression and interferon-gamma response in melanocytic cells. Pigm Cell Melanoma R 31, 51–63 (2018).

34.) Wang, C., et al. Donkey genomes provide new insights into domestication and selection for coat color. Nat Commun 11, 6014 (2020).

35.) Imsland, F., et al. Regulatory mutations in TBX3 disrupt asymmetric hair pigmentation that underlies Dun camouflage color in horses. Nat Genet 48, 152–158 (2016).

36.) Yin, K., Sturm, R. A. & Smith, A. G. MC1R and NR4A receptors in cellular stress and DNA repair: implications for UVR protection. Exp Dermatol 23, 449–452 (2014).

37.) Moon, H., et al. Melanocyte Stem Cell Activation and Translocation Initiate Cutaneous Melanoma in Response to UV Exposure. Cell Stem Cell 21, 665–678.e6 (2017).

38.) Zhang, T., et al. Cell-type–specific eQTL of primary melanocytes facilitates identification of melanoma susceptibility genes. Genome Res 28, 1621–1635 (2018).

39.) Perotti, V., et al. NFATc2 is an intrinsic regulator of melanoma dedifferentiation. Oncogene 35, 2862–2872 (2016).

40.) Smith, A. G., Lim, W., Pearen, M., Muscat, G. E. O. & Sturm, R. A. Regulation of NR4A nuclear receptor expression by oncogenic BRAF in melanoma cells. Pigm Cell Melanoma R 24, 551–563 (2011).

41.) Boyd, S. C., et al. Oncogenic B-RAFV600E Signaling Induces the T-Box3 Transcriptional Repressor to Repress E-Cadherin and Enhance Melanoma Cell Invasion. J Invest Dermatol 133, 1269–1277 (2013).

42.) Lu, J., Li, X.-P., Dong, Q., Kung, H. & He, M.-L. TBX2 and TBX3: The special value for anticancer drug targets. Biochimica Et Biophysica Acta Bba - Rev Cancer 1806, 268–274 (2010).

43.) Peres, J. et al. TBX3 promotes melanoma migration by transcriptional activation of ID1 which prevents activation of E-cadherin by MITF. J Invest Dermatol (2021) doi:10.1016/j.jid.2021.02.740.

44.) Rodriguez, M., Aladowicz, E., Lanfrancone, L. & Goding, C. R. Tbx3 Represses E-Cadherin Expression and Enhances Melanoma Invasiveness. Cancer Res 68, 7872– 7881 (2008).

45.) Peres, J., et al. The Highly Homologous T-Box Transcription Factors, TBX2 and TBX3, Have Distinct Roles in the Oncogenic Process. Genes Cancer 1, 272–282 (2010).

46.) Perrot, C. Y., et al. GLI2 cooperates with ZEB1 for transcriptional repression of CDH1 expression in human melanoma cells. Pigm Cell Melanoma R 26, 861–873 (2013).

47.) Heppt, M. V., et al. MSX1-Induced Neural Crest-Like Reprogramming Promotes Melanoma Progression. J Invest Dermatol 138, 141–149 (2018).

48.) Chen, Y., et al. ZEB1 Regulates Multiple Oncogenic Components Involved in Uveal Melanoma Progression. Sci Rep-uk 7, 45 (2017).

49.) Goding, C. R. & Arnheiter, H. MITF—the first 25 years. Gene Dev 33, 983–1007 (2019).

50.) Murakami, M., Matsuzaki, F. & Funaba, M. Regulation of melanin synthesis by the TGF-β family in B16 melanoma cells. Mol Biol Rep 36, 1247–1250 (2009).

51.) Li, J., Weinberg, M. S., Zerbini, L. & Prince, S. The oncogenic TBX3 is a downstream target and mediator of the TGF-β1 signaling pathway. Mol Biol Cell 24, 3569–3576 (2013).

52.) Laurette, P., et al. Transcription factor MITF and remodeller BRG1 define chromatin organisation at regulatory elements in melanoma cells. Elife 4, e06857 (2015).

53.) Louphrasitthiphol, P., et al. Tuning Transcription Factor Availability through Acetylation-Mediated Genomic Redistribution. Mol Cell 79, 472–487.e10 (2020).

54.) Lu, S., et al. TBX2 controls a proproliferative gene expression program in melanoma. Gene Dev 35, 1657–1677 (2021).

55.) McLean, C. Y., et al. GREAT improves functional interpretation of cis-regulatory regions. Nat Biotechnol 28, 495–501 (2010).

56.) Strub, T., et al. Essential role of microphthalmia transcription factor for DNA replication, mitosis and genomic stability in melanoma. Oncogene 30, 2319–2332 (2011).

57.) Praetorius, C., et al. A Polymorphism in IRF4 Affects Human Pigmentation through a Tyrosinase-Dependent MITF/TFAP2A Pathway. Cell 155, 1022–1033 (2013).

58.) Aoki, H. & Moro, O. Involvement of microphthalmia-associated transcription factor (MITF) in expression of human melanocortin-1 receptor (MC1R). Life Sci 71, 2171–2179 (2002).

59.) Visser, M., Kayser, M. & Palstra, R.-J. HERC2 rs12913832 modulates human pigmentation by attenuating chromatin-loop formation between a long-range enhancer and the OCA2 promoter. Genome Res 22, 446–455 (2012).

60.) Gautron, A. et al. Human TYRP1: two functions for a single gene? Pigm Cell Melanoma R (2020) doi:10.1111/pcmr.12951.

61.) Washkowitz, A. J., Gavrilov, S., Begum, S. & Papaioannou, V. E. Diverse functional networks of Tbx3 in development and disease. Wiley Interdiscip Rev Syst Biology Medicine 4, 273–283 (2012).

62.) Khan, S. F., et al. Gene 726, 144223 (2019).

63.) Pape, E. L., Wakamatsu, K., Ito, S., Wolber, R. & Hearing, V. J. Regulation of eumelanin/pheomelanin synthesis and visible pigmentation in melanocytes by ligands of the melanocortin 1 receptor. Pigm Cell Melanoma R 21, 477–486 (2008).

64.) Ni, J., et al. The effect of the NMDA receptor-dependent signaling pathway on cell morphology and melanosome transfer in melanocytes. J Dermatol Sci 84, 296–304 (2016).

65.) Tagore, M. et al. Electrical activity between skin cells regulates melanoma initiation. Biorxiv 2021.12.19.473393 (2021) doi:10.1101/2021.12.19.473393.

66.) Enkhtaivan, E. & Lee, C. H. Role of Amine Neurotransmitters and Their Receptors in Skin Pigmentation: Therapeutic Implication. Int J Mol Sci 22, 8071 (2021).

67.) Palumbo, A. Melanogenesis in the Ink Gland of Sepia officinalis. Pigm Cell Res 16, 517– 522 (2003).

68.) Eddy, K. & Chen, S. Glutamatergic Signaling a Therapeutic Vulnerability in Melanoma. Cancers 13, 3874 (2021).

69.) Janes, P. W., et al. EphA3 biology and cancer. Growth Factors 32, 176–189 (2014).

70.) Theveneau, E. & Mayor, R. Neural crest migration: interplay between chemorepellents, chemoattractants, contact inhibition, epithelial–mesenchymal transition, and collective cell migration. Wiley Interdiscip Rev Dev Biology 1, 435–445 (2012).

71.) Pape, E. L., et al. Microarray analysis sheds light on the dedifferentiating role of agouti signal protein in murine melanocytes via the Mc1r. Proc National Acad Sci 106, 1802– 1807 (2009).

72.) Arnheiter, H. The discovery of the microphthalmia locus and its gene, Mitf. Pigm Cell Melanoma R 23, 729–735 (2010).

73.) Levy, C., Khaled, M. & Fisher, D. E. MITF: master regulator of melanocyte development and melanoma oncogene. Trends Mol Med 12, 406–414 (2006).

74.) Li, J., et al. The Anti-proliferative Function of the TGF-β1 Signaling Pathway Involves the Repression of the Oncogenic TBX2 by Its Homologue TBX3*. J Biol Chem 289, 35633– 35643 (2014).

75.) Mesbah, K., et al. Identification of a Tbx1/Tbx2/Tbx3 genetic pathway governing pharyngeal and arterial pole morphogenesis. Hum Mol Genet 21, 1217–1229 (2012).

76.) Mohan, R. A., et al. T-box transcription factor 3 governs a transcriptional program for the function of the mouse atrioventricular conduction system. Proc National Acad Sci 117, 18617–18626 (2020).

77.) Singh, R., et al. Tbx2 and Tbx3 induce atrioventricular myocardial development and endocardial cushion formation. Cell Mol Life Sci 69, 1377–1389 (2012).

78.) Lüdtke, T. H., et al. Tbx2 and Tbx3 Act Downstream of Shh to Maintain Canonical Wnt Signaling during Branching Morphogenesis of the Murine Lung. Dev Cell 39, 239–253 (2016).

79.) Sheeba, C. J. & Logan, M. P. O. Chapter Twelve The Roles of T-Box Genes in Vertebrate Limb Development. Curr Top Dev Biol 122, 355–381 (2017).

80.) Frank, D. U., Emechebe, U., Thomas, K. R. & Moon, A. M. Mouse Tbx3 Mutants Suggest Novel Molecular Mechanisms for Ulnar-Mammary Syndrome. Plos One 8, e67841 (2013).

81.) Kaiser, M., et al. Regulation of otocyst patterning by Tbx2 and Tbx3 is required for inner ear morphogenesis in the mouse. Development 148, dev.195651 (2021).

82.) Dong, L., et al. Novel HDAC5-interacting motifs of Tbx3 are essential for the suppression of E-cadherin expression and for the promotion of metastasis in hepatocellular carcinoma. Signal Transduct Target Ther 3, 22 (2018).

83.) Peres, J. & Prince, S. The T-box transcription factor, TBX3, is sufficient to promote melanoma formation and invasion. Mol Cancer 12, 117 (2013).

84.) Hoekstra, H. E. Genetics, development and evolution of adaptive pigmentation in vertebrates. Heredity 97, 222–234 (2006).

85.) Chintala, S., et al. Slc7a11 gene controls production of pheomelanin pigment and proliferation of cultured cells. P Natl Acad Sci Usa 102, 10964–10969 (2005).

86.) Poltorack, C. D. & Dixon, S. J. Understanding the role of cysteine in ferroptosis: progress & paradoxes. Febs J 289, 374–385 (2022).

87.) Willmer, T., Peres, J., Mowla, S., Abrahams, A. & Prince, S. The T-Box factor TBX3 is important in S-phase and is regulated by c-Myc and cyclin A-CDK2. Cell Cycle 14, 3173– 3183 (2015).

88.) Damerell, V., et al. The c-Myc/TBX3 Axis Promotes Cellular Transformation of Sarcoma-Initiating Cells. Frontiers Oncol 11, 801691 (2022).

89.) Coll, M., Seidman, J. G. & Müller, C. W. Structure of the DNA-Bound T-Box Domain of Human TBX3, a Transcription Factor Responsible for Ulnar-Mammary Syndrome. Structure 10, 343–356 (2002).

90.) Langmead, B. & Salzberg, S. L. Fast gapped-read alignment with Bowtie 2. Nat Methods 9, 357–359 (2012).

91.) Zhang, Y., et al. Model-based Analysis of ChIP-Seq (MACS). Genome Biol 9, R137 (2008).

92.) Bailey, T. L., Johnson, J., Grant, C. E. & Noble, W. S. The MEME Suite. Nucleic Acids Res 43, W39–W49 (2015).

93.) Quinlan, A. R. & Hall, I. M. BEDTools: a flexible suite of utilities for comparing genomic features. Bioinformatics 26, 841–842 (2010).

94.) Yu, G., Wang, L.-G. & He, Q.-Y. ChIPseeker: an R/Bioconductor package for ChIP peak annotation, comparison and visualization. Bioinformatics 31, 2382–2383 (2015).

