## Supplemental Figures for "Loss of MC1R signaling implicates TBX3 in pheomelanogenesis and melanoma predisposition"

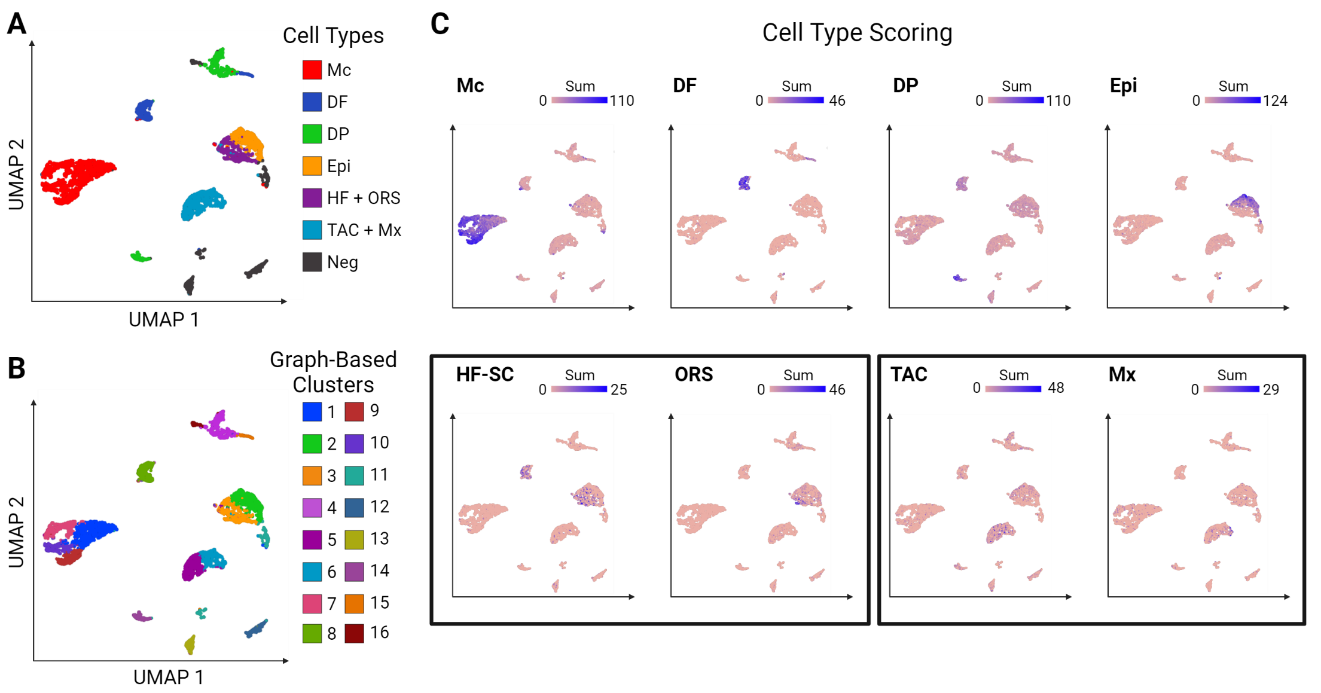

**Fig S1. Classification Scheme for nonagouti and lethal yellow cells.** (A) Uniform Manifold Approximation and Projection (UMAP) plot of lethal yellow and nonagouti cells classified by skin cell type: Melanocytes (Mc), Dermal fibroblasts (DF), Dermal papilla (DP), Epidermis (Epi), Hair follicle and Outer root sheath (HF + ORS), and Transit amplifying and Matrix cells (TAC + Mx). (B) UMAP of cells classified by graph-based cluster. (C) UMAPs depicting heatmap of cell type scoring.

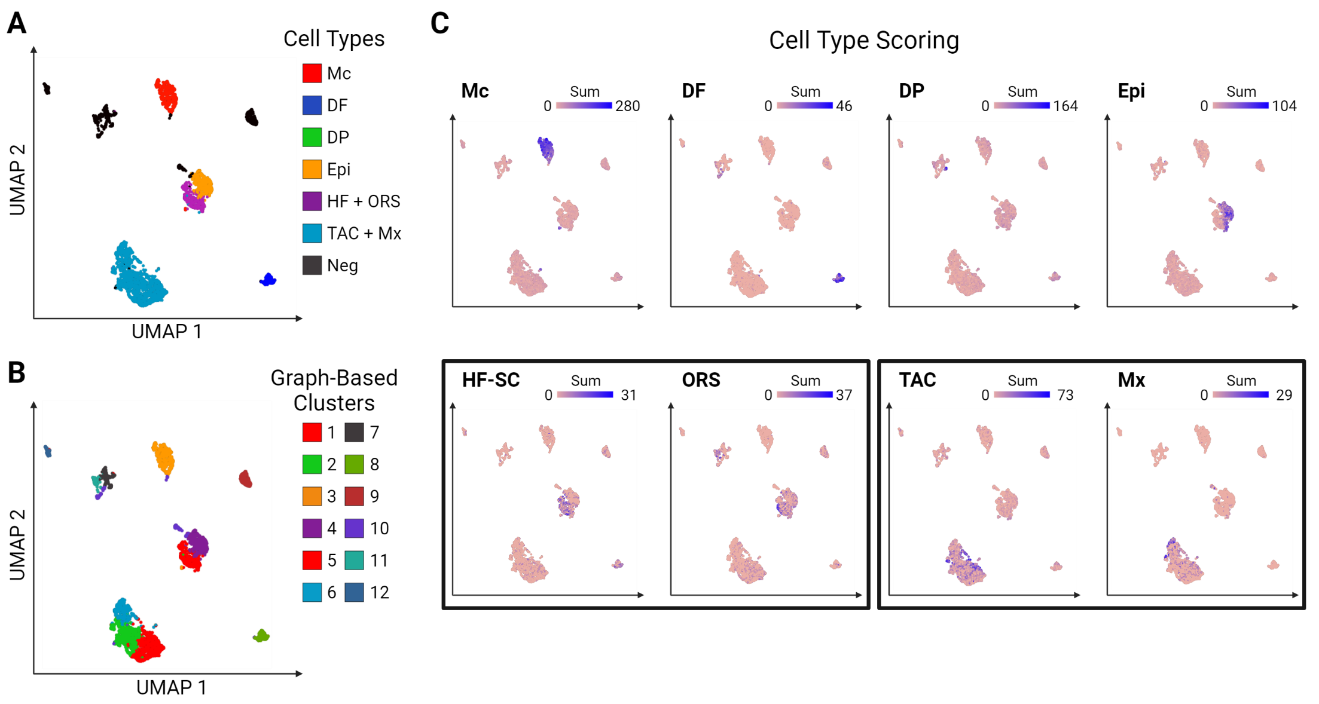

**Fig S2. Classification Scheme for nonagouti and recessive yellow cells.** (A) Uniform Manifold Approximation and Projection (UMAP) plot of the recessive yellow and nonagouti cells classified by skin cell type: Melanocytes (Mc), Dermal fibroblasts (DF), Dermal papilla (DP), Epidermis (Epi), Hair follicle and Outer root sheath (HF + ORS), and Transit amplifying and Matrix cells (TAC + Mx). (B) UMAP of cells classified by graph-based cluster. (C) UMAPs depicting heatmap of cell type scoring.

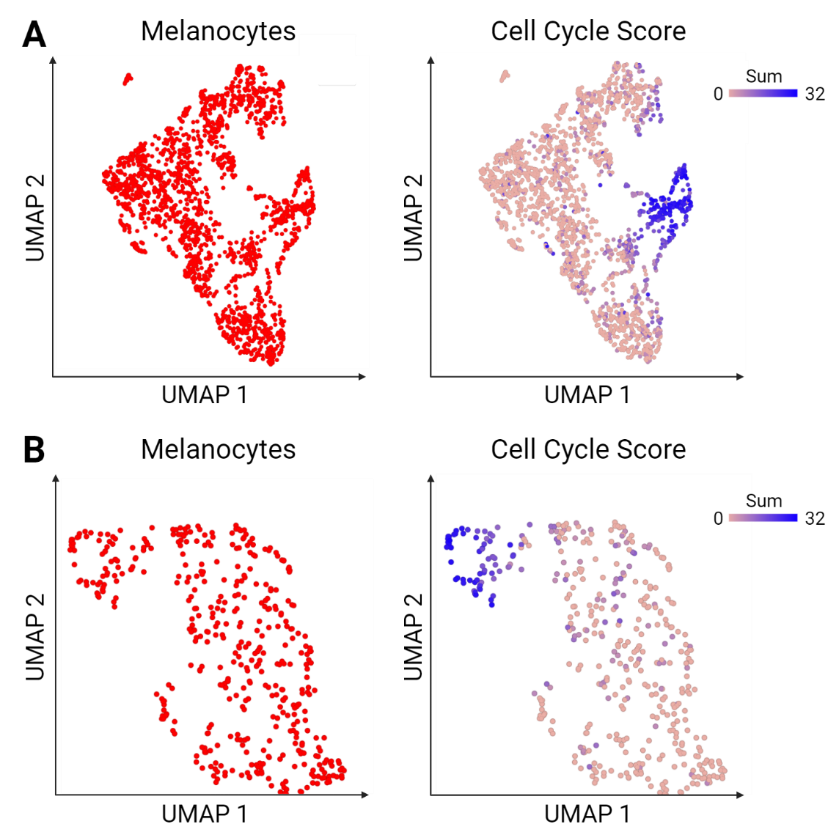

**Fig S3. Melanocytes cell cycle.** (A) Total melanocytes from lethal yellow and nonagouti pairwise experiment (left), cells within the cell cycle highlighted in blue (right). (B) Total melanocytes from recessive yellow and nonagouti pairwise experiment (left), cells within the cell cycle highlighted in blue (right).

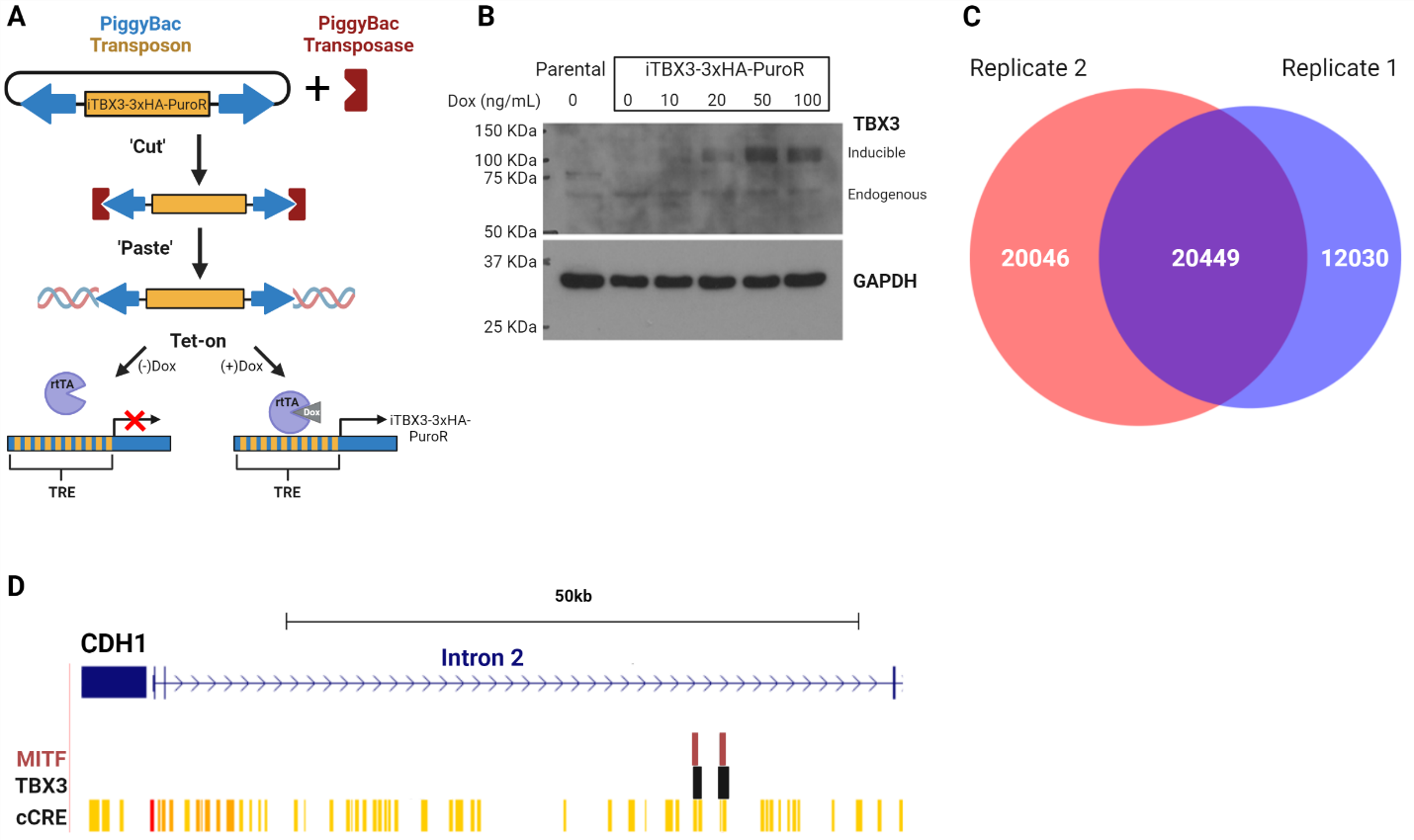

**Fig S4. Inducible TBX3 cell line validation.** (A) schematic of piggyBAC transposase system with Tet-on inducible expression of TBX3. Dox, doxycycline; TRE, tetracycline response element; rtTA, reverse tetracycline-controlled transactivator. (B) Western blot of polyclonal 501mel-iTBX3-3xHA-PuroR inducible cells treated with varying amounts of doxycycline for 24h. (C) Venn diagram depicting shared peaks between both replicate ChIP experiments. (D) UCSC browser view of TBX3 and MITF ChIP peaks associated with E-cadherin. Browser tracks are as follows: TBX3 = replicated binding sites in our dataset; MITF = ChIP peaks from Strub et al; cCRE=encode annotated candidate cis-regulatory elements.

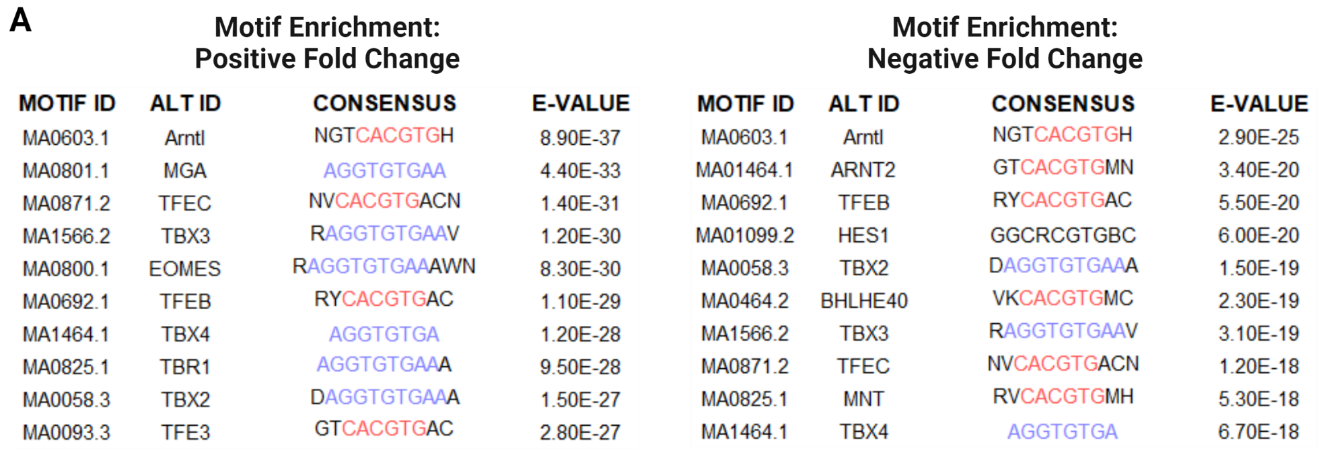

**Figure S5. Motif enrichment by fold-change direction in TBX3 KD studies.** (A) Top 10 motifs, ranked by e-value, of TBX3 ChIP peaks associated with putative target genes with increased expression (left) or decreased expression (right) upon TBX3 knock-down.
